# Predicting the protein interaction landscape of a free-living bacterium with pooled-AlphaFold3

**DOI:** 10.1101/2025.07.01.662654

**Authors:** Horia Todor, Lili M. Kim, Jürgen Jänes, Hannah N. Burkhart, Seth A. Darst, Pedro Beltrao, Carol A. Gross

## Abstract

Accurate prediction of protein complex structures by AlphaFold3 and similar programs has been used to predict the presence of protein-protein interactions (PPIs), but this technique has never been applied to an entire genome due to onerous computational requirements and questionable utility. Here we present pooled-PPI prediction, a technique that dramatically improves the accuracy of genome-scale screens compared to a paired approach while simultaneously reducing inference time (∼2-fold) and the number of jobs (∼100-fold). We use this technique to predict the structure of all 113,050 pairwise PPIs in *Mycoplasma genitalium* using only 2,027 AlphaFold3 jobs. This unbiased and comprehensive dataset was highly predictive of known interactions, revealed a previously unappreciated but widespread size bias in AlphaFold interface scores, correctly identified protein-protein interfaces in macromolecular complexes, and uncovered new biology in *M. genitalium.* This work establishes pooled-PPI prediction as a highly scalable method for uncovering protein-protein interactions and a powerful addition to the functional genomics toolkit.

## Main Text

Deciphering the function of novel proteins based on their primary sequence is a longstanding goal in biology. The release of AlphaFold2 in 2021 represented a significant step towards this goal, predicting protein structures with unprecedented accuracy (*1*). However, discerning the function of a novel protein, even knowing its structure, remains a significant challenge. Both AlphaFold2 and its successor AlphaFold3 are capable of predicting the structures of protein complexes and assigning a score to predicted protein-protein interfaces in such complexes (*2*, *3*). Associating an unannotated protein with a well-understood partner is an important and informative step to deciphering its function. Although both AlphaFold 2 and AlphaFold3 were designed to predict *how* proteins interact, both have been successfully co-opted to predict *whether* two proteins interact, and an increasingly large body of evidence points to the utility of genome-scale protein-protein interaction (PPI) prediction profiling for dissecting biological processes and determining gene function (*4–8*). However, the use of genome-scale PPI prediction is limited by its onerous computational requirements and the potential for false positive hits. While there has been a significant effort to make protein structure prediction programs accessible to non-expert users (*9*) (such as the web interfaces for AlphaFold3 and CHAI1 and the ColabFold notebook for AlphaFold2 and AlphaFold-Multimer), the limited nature of these interfaces make them unsuitable for genome-scale predictions.

Here, we describe and characterize pooled-PPI prediction, a broadly applicable strategy for facilitating genome-scale screens. Relying on the fact that PPIs are rare (e.g. ∼0.05% of possible protein pairs are thought to interact in humans) (*10*), pooled-PPI prediction assumes that predicted structure (and therefore interface score) of a true complex is unlikely to be affected by the addition of other, non-interacting proteins in the same job. Including multiple proteins in a single job allows numerous potential pairwise interactions (*n* choose 2) to be screened simultaneously. Although mutually exclusive interactions sharing the same interface could be missed if multiple partners are in the same pool, the chance of multiple binding partners being in the same pool is low in a genome-wide approach. Pooled-PPI prediction decreases overall run-time (∼2-fold, depending on hardware), the number of individual jobs (up to 300-fold, depending on pool size), and increases the accuracy of PPI predictions because of its intrinsically competitive nature (*11–13*).

We demonstrate the utility of this method by performing a comprehensive genome-wide (all by all) predicted PPI screen for the free-living bacterium *Mycoplasma genitalium* (113,050 unique protein pairs) using only the free AlphaFold3 web interface. We make several findings. First and most importantly, we found that pooled-AlphaFold3 predictions recapitulate experimentally validated PPIs in *M. genitalium* while drastically reducing the number of false positives compared to pairwise predictions. Second, the scale and unbiased nature of our screen revealed a widespread size bias in AlphaFold3 interaction scores which is not due to our pooled approach. Correcting for the size bias not only drastically improved the identification of biologically relevant interactions but also revealed significant variability in AlphaFold3 interface scores in both paired and pooled contexts. Third, we found that despite assessing interactions in a pairwise manner, we could reconstruct macromolecular machines such as the ribosome and RNA polymerase. Finally, we discovered novel PPIs in *M. genitalium,* including many with orthogonal experimental support, that suggest new biology. Together, these results establish pooled-PPI prediction as a powerful and broadly applicable method for facilitating genome-scale PPI screens.

## Results

### Pooled-PPI prediction improves runtime and decreases the total number of runs

Pooled-PPI prediction leverages the known rarity of PPIs (*10*) to evaluate multiple potential PPIs in a single job by co-folding several proteins rather than only two (Fig. 1A). Since the number of potential pairwise PPIs predicted in a single job scales quadratically (as *n* choose 2) with the number of proteins (Fig. 1B), this strategy can drastically reduce the number of jobs required to predict a given set of PPIs. The number of proteins that can be included in a job depends on the protein size and the prediction algorithm. The web interface for AlphaFold3 accepts inputs up to 5,000 tokens (for PPI prediction, 5,000aa) (*3*). However, larger jobs require more computational time (*3*, *14*) (Fig. 1C), and this scaling appears to be approximately quadratic for AlphaFold3. To determine the theoretical benefit of pooled-PPI, we considered hypothetical jobs with proteins of 200aa to 500aa - a range that encompasses both the median bacterial protein (∼300aa) and the median human protein (∼480aa) - and estimated the runtime of a paired and pooled approach. We found that pooling decreases total inference time by 1.4- to 2-fold, depending on the underlying hardware (Fig. 1D). Intuitively, this performance benefit arises from the fact that in a comprehensive pooled approach each individual protein is folded less often than in a comparable paired approach. Importantly, the total number of jobs required decreases significantly (45- to 300-fold), allowing many more PPIs to be assayed using the free allotment of jobs provided by online resources such as the AlphaFold3 web interface (currently 30 jobs per day per user). The theoretical benefits of pooling are even more significant for higher order interactions - a single job with 10 proteins assesses 120 tripartite interactions while one with 25 assesses 2,300 (Fig. S1), resulting in a 16- to 25-fold improvement in overall runtime for querying such interactions.

**Fig. 1.**
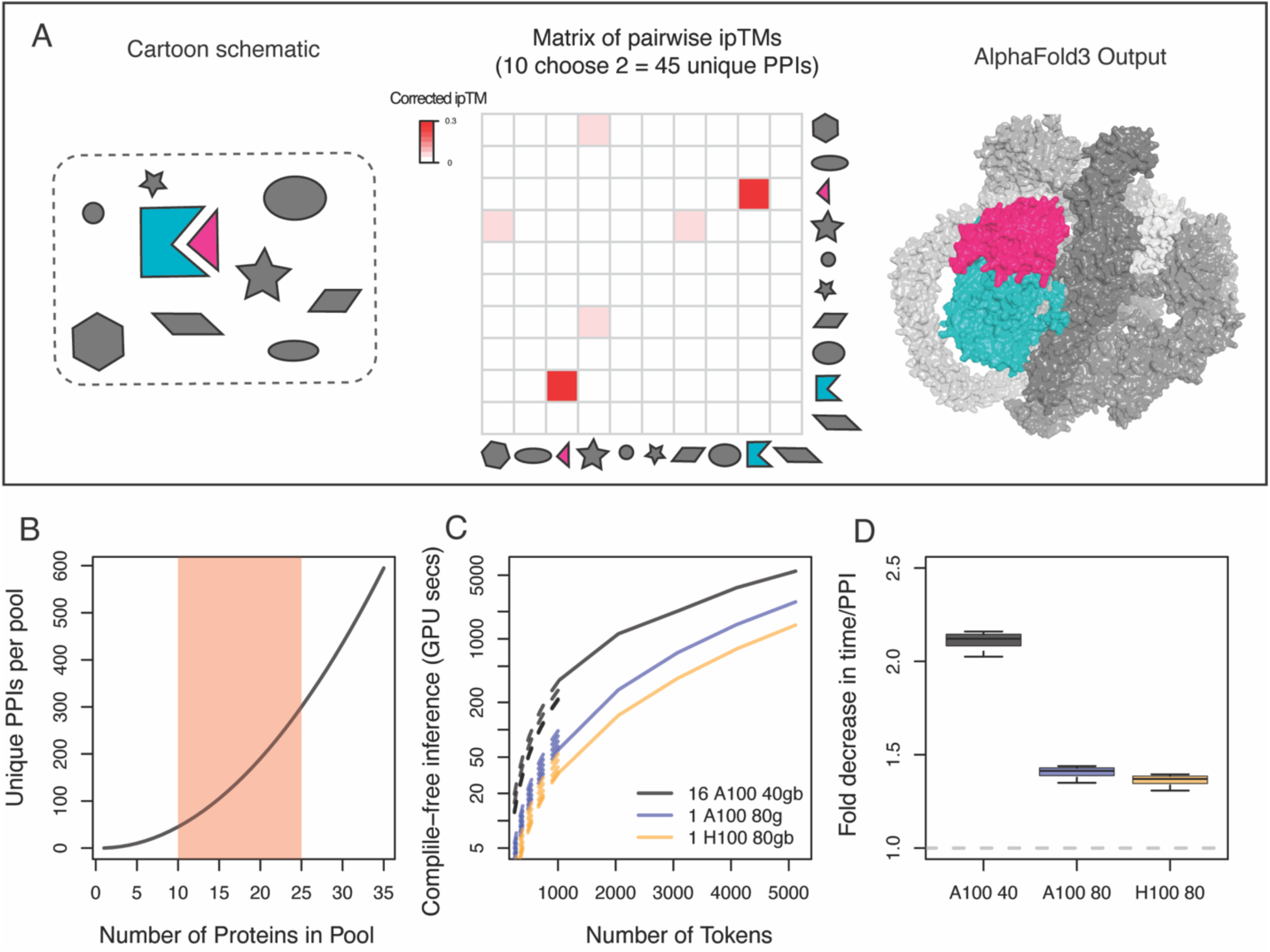
Pooled-PPI prediction reduces the number of jobs and increases inference throughput. **A.** Schematic describing pooled-PPI. A job (250310_mgen_allbyall_627) containing 10 proteins produces a matrix of size-corrected pairwise ipTMs. 45 unique potential interactions are assayed in this job. Pooled-AlphaFold3 correctly identifies an interacting protein pair in the milieu of non-interacting pairs (MG_290 (blue) and MG_291 (red), ABC-transporter components). **B.** The number of proteins in the pool determines the number of unique pairwise interactions screened in a single job. The red shaded area encompasses most jobs. **C.** AlphaFold3 compile-free inference time increases as the job size increases in a GPU-dependent function. Broken lines indicate estimates from each published timing (1024-5120 tokens) based on the assumption of quadratic scaling, since the runtime for jobs <1,024 tokens are not published. **D.** Pooled-PPI-prediction decreases inference time per unique PPI even considering the increased runtime of larger jobs. All timing estimates (1024-5120 tokens) and pool sizes of 10-25 proteins are considered and reflected in the error bars.

### Prediction of all pairwise PPIs in a free-living organism reveals size bias in AlphaFold3 interface scores

To characterize the performance of the pooled-PPI approach, we used it to predict all pairwise PPIs in the free-living bacterium *M. genitalium* (476 proteins, 113,050 unique protein pairs, Table S2) with AlphaFold3. Each of the 2,027 jobs contained a random selection of 4-23 proteins whose sizes sum to almost 5,000 amino acids (median *n* = 13, Fig. S2, Table S2, Methods), and together queried a total of 160,772 total pairwise PPIs (some protein pairs appear in more than one pool) as well as 636,516 unique tripartite PPIs (3.56% of 17,861,900 such interactions). These jobs took 68 person-days to run using the free allotment of jobs on the AlphaFold3 web interface.

AlphaFold3 evaluates the accuracy of predicted complexes using the interface predicted template modeling (ipTM) score (*2*, *3*). Generally, ipTM values >0.8 are considered confident predictions, those between 0.6 and 0.8 are a grey area, and those <0.6 are incorrect predictions (*2*, *3*). Consistent with the expectation that protein interactions are rare, very few protein pairs had high ipTM scores: 98.9% were between 0 and 0.2 (Fig. 2A).

**Fig. 2.**
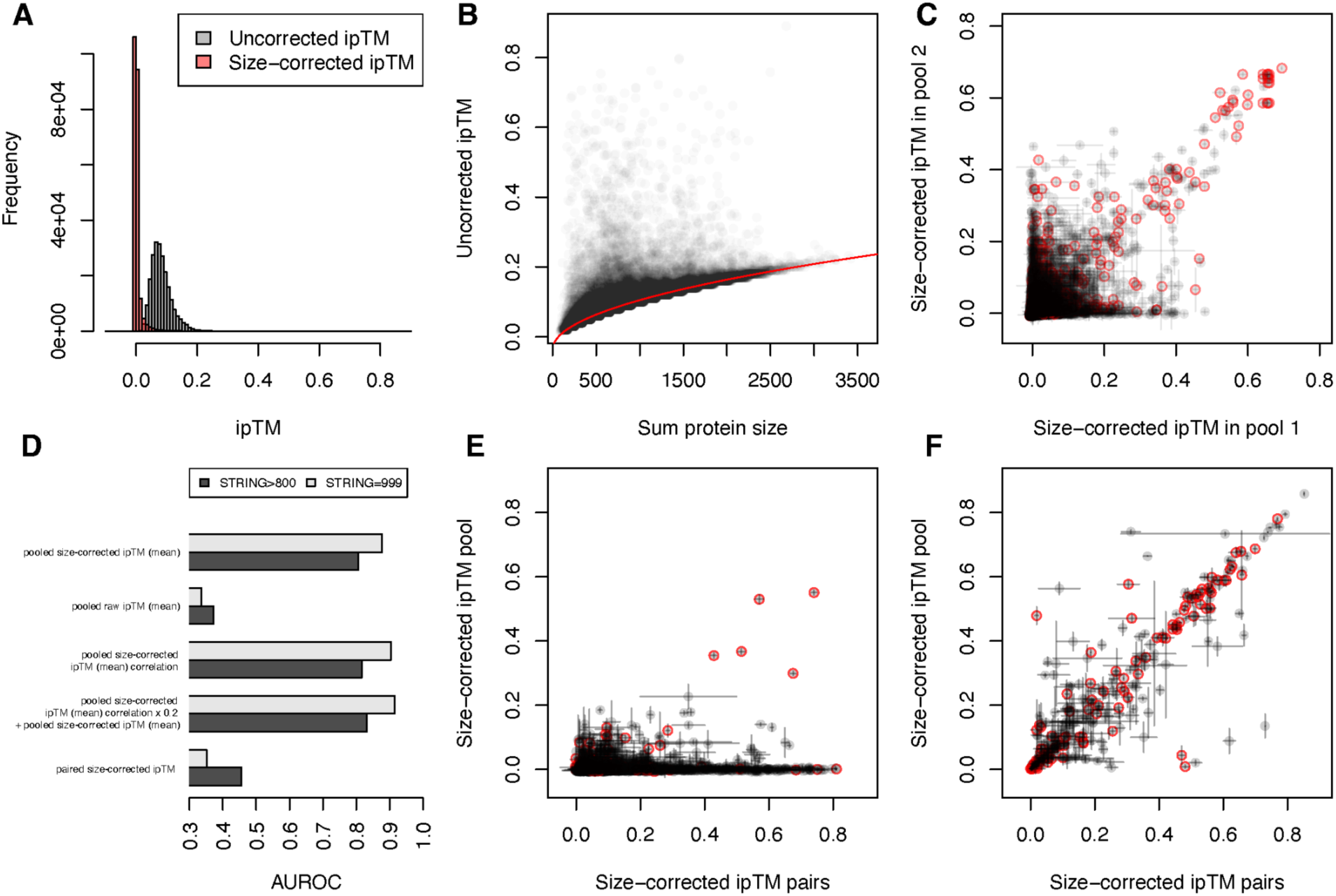
Technical characterization of pooled-AlphaFold3. **A.** Distribution of raw (grey) and corrected (red) pairwise ipTM scores for all 113,050 unique protein pairs in the *M. genitalium* genome. **B.** The ipTM for non-interacting protein pairs is proportional to the square root of the summed size of the proteins. The red line is sqrt(sum_protein_size)*0.0044 - 0.036. **C.** Pooled-AlphaFold3 scores are reproducible. Size-corrected ipTM scores for the 38,718 protein pairs that appear in multiple pools are similar (Pearson’s *r* = 0.670, 38,718 protein pairs). **D.** Pooled-AlphaFold3 accurately predicts known interactions in the STRING database. AUROC scores for different variations compared to the STRING experimental database. **E.** Pooled-AlphaFold3 scores are poorly correlated with paired approaches. Size-corrected ipTM differs between paired and pooled AlphaFold3 jobs (Pearson’s *r* = 0.156, 4,560 random pairs). **F.** AlphaFold3 has intrinsic variability in ipTM scores. Size-corrected ipTM scores across replicated paired runs exhibit surprising variability (Pearson’s *r* = 0.878, 314 pairs). For **C, E,** and **F**, circles in red are interactions with scores > 800 in the experimental channel of the STRING database.

The relative paucity of protein pairs with ipTM ∼0.0 and substantial width of the ipTM distribution (Fig. 2A) for putatively non-interacting pairs was surprising, considering that many of those protein pairs were modeled >30 Å apart. This observation led us to wonder whether some aspect of protein pairs other than their interface may influence their ipTM. Analysis revealed a strong and significant correlation between ipTM and the square root of the summed size of the interacting proteins (robust R^2^ = 0.983, Fig. 2B, Methods). Since there is no apparent biological reason for this size-ipTM relationship, we predicted ipTM for each pair based on the sum of the protein sizes and subtracted this prediction from all of our ipTM values to generate size-corrected ipTM scores (Methods, Supplementary Text 1). Size-corrected ipTM exhibited a much narrower distribution of scores for non-interacting protein pairs (Fig. 2A), and (as discussed below) were more predictive of biologically relevant interactions. A similar size bias appears in published AlphaFold-multimer datasets (Fig. S3A-B) and in our AlphaFold3 predictions of protein pairs (Fig. S3C), suggesting that this bias is not due to our pooled approach but represents a more general (and potentially undesirable) feature of ipTM scores.

### Pooled-AlphaFold3 is reproducible

To assess the reproducibility of pooled ipTM measurements, we first analyzed the 38,718 protein pairs that were present in 2 or more pools. The size-corrected ipTM scores of these 38,718 proteins were well correlated in the different pools (Pearson’s *r* = 0.670, Fig. 2C), especially for pairs with a size-corrected ipTM > 0.4, suggesting minimal context dependence.

Performing the same analysis using raw ipTM scores gave the illusion of better reproducibility (Pearson’s *r* = 0.83, Fig. S4, Table S2), because the effect of size on raw ipTM is the same across replicates. These data validate our assumption that pool context does not strongly affect the outcome of PPI predictions.

### M. genitalium predicted PPIs recapitulate known interactions

Having validated the technical aspects of our screen, we next tested how well the predicted PPIs captured known biological interactions. *Mycoplasma* species (like *M. genitalium*) are ideal test cases: they have streamlined genomes and have been well studied from clinical (*15*), synthetic (*16*), and basic (*17*, *18*) biology perspectives. We benchmarked our *M. genitalium* predicted-PPI dataset against the STRING database “experimental” channel, which consists of protein–protein association evidence imported from repositories such as BioGRID, PDB, and IntAct rigorously propagated through bacterial phylogeny to appropriately integrate information from related species (*19*). As expected considering the known rarity of PPIs (*10*), the vast majority (105,338/113,050, 93.2%) of protein pairs in *M. genitalium* had no evidence of interaction in the STRING experimental channel. Only 2,633 protein pairs (2.3%) had strong evidence of interaction (score >800), and even fewer (1,487, 1.3%) had the strongest evidence of interaction (score = 999). The average number of strong interactions per protein (5.53 in *M. genitalium*) is consistent with that of essential genes in well studied bacteria (e.g. 4.12 strong interactions per essential protein in *Bacillus subtilis*), suggesting that STRING adequately captures known PPIs in *M. genitalium.* Size-corrected ipTM scores were predictive of these known interactions, as quantified by the area under the receiver operating characteristic curves (AUROC), exhibiting an AUROC of 0.81 for strong interactions and 0.88 for the strongest interactions (Fig. 2D, Fig. S5, Table S4). Using the raw (not size corrected) ipTM values performed significantly worse (Fig. S5, AUROC = 0.37 and 0.34 for strong and the strongest interactions, respectively), supporting our correction of this bias and raising the possibility that applying this correction could reveal new biologically relevant interactions in other data sets.

The worse than random AUROC of raw ipTMs is due to the fact that interacting protein pairs are generally shorter than random pairs (median protein pair lengths: strong evidence of interaction 346aa, random 655aa). Strikingly, we found that the predictive power of our size-corrected ipTM scores was not limited solely to the strongest interactions (Table S4): protein pairs with size-corrected ipTM 0.1-0.2 were also significantly enriched in strong (5.8-fold) and the strongest (8.2-fold) STRING experimental interactions, as were protein pairs with size-corrected ipTM 0.05-0.1 (4.8- and 6.4-fold, respectively).

We also tested whether correlations between the predicted PPI profiles of different proteins can indicate additional functional relationships, as would be the case if two otherwise non-interacting proteins both interact with the same partner(s). Consistent with this interpretation, the correlation matrix of predicted PPIs (Table S4) was also predictive of both strong (AUROC = 0.82) and the strongest (AUROC = 0.90) interactions in the STRING experimental database (Fig. 2D, Fig. S5). While protein pairs with high size-corrected ipTM were more likely to have strongly correlated partners, numerous protein pairs without strong evidence of interaction also exhibited correlated PPI profiles (Fig. S6A), suggesting that the correlation of PPI profiles represents a partially orthogonal source of information that can potentially overcome defects in PPI prediction. To test this hypothesis, we combined the size-corrected ipTM matrix with the correlation matrix in varying proportions and assessed the performance of these combined scores in predicting interactions in the STRING experimental database. Combined scores slightly outperformed both individual metrics (Fig. 2D); a combined score consisting of size-corrected ipTM + 0.2 * correlation achieved good predictions for both strong (AUROC = 0.83) and for the strongest interactions (AUROC = 0.92, Fig. 2D, Fig. S4, Fig. S6B). Together, these data demonstrate that pooled-AlphaFold3 sensitively and specifically captures known PPIs.

#### Pooled-AlphaFold3 outperforms a paired approach for predicting PPIs

Previous *in silico* PPI screens were performed by co-folding two proteins per job (*4–8*). To compare pairwise predictions to our pooled approach, we used AlphaFold3 to individually co-fold a representative set of 4,560/113,050 unique *M. genitalium* protein pairs (Methods, Table S3) and ascertained their ability to predict known PPIs. Surprisingly, the size-corrected ipTM of individually co-folded pairs was substantially worse than our pooled approach at identifying both strong (AUROC = 0.46 compared to 0.81) and the strongest (AUROC = 0.35 compared to 0.88) interactions in the STRING experimental database (Fig. 2D, Table S3). Consistent with this, size-corrected ipTMs of protein pairs were poorly correlated between the paired and pooled approaches (Pearson’s *r* = 0.157), driven primarily by numerous protein pairs with significant size-corrected ipTM when folded individually and near zero size-corrected ipTM in pools (Fig. 2E). We additionally individually co-folded 942 protein pairs covering most of the high size-corrected ipTM interactions identified by the pooled approach. This set of protein pairs exhibited highly correlated size-corrected ipTM between the pooled and paired contexts, (Pearson’s r = 0.725, Fig S7A), suggesting a low false-positive rate for the pooled approach vis-a-vis individually co-folded protein pairs.

That pooled-AlphaFold3 outperforms a paired approach is consistent with previous work showing that “competitive” AlphaFold protein-peptide binding can improve predictions (*11–13*). Since these papers used older versions of AlphaFold and tested only small numbers (i.e. 2 to 5) of competitive peptides per job, we next asked whether 5,000aa pools are required to achieve the full benefits of pooling. To answer this question, we randomly partitioned all *M. genitalium* proteins (n=476) into 5 approximately equal groups, then modelled all interactions within a group using either pairs or random pools of increasing size (2,000aa, 3,000aa, 4,000aa, or 5,000aa) and assessed how well their size-corrected ipTM predicted known interactions (STRING experimental > 800). Individually folded pairs had the lowest AUROC, and AUROC steadily increased as pool size increased up to 5,000aa (Fig. S7B-C, Table S5), suggesting that the full benefit of pooling requires at least 5,000aa pools. The increased AUROC was driven primarily by a decrease in false-positive hits (Fig. S7D-F). We were unable to test larger pool sizes due to a lack of GPUs with >>80gb of memory, but it is possible that larger pools could improve performance further. Our results significantly extend previous observations on the benefits of pooling (*11–13*) and suggest that large pools with many proteins dramatically enhance the utility of genome-scale PPI predictions by reducing false positive interactions.

### Averaging inherently variable AlphaFold3 interface scores modestly improves PPI prediction

When comparing the size-corrected ipTM across pools, much of the variability arose from protein pairs that exhibit a meaningful size-corrected ipTM (0.2-0.4) in one run, and near zero size-corrected ipTM in another. This observation led us to ask whether the variability was due to our pooled approach or if it is inherent to the generative framework of AlphaFold3. To determine the source of the variability, we ran 314 individual pairs twice using different random seeds.

Strikingly, these individually folded pairs also exhibited substantial variability in size-corrected ipTM (Pearson’s *r* = 0.878, Fig. 2F). Of the 314 protein pairs that were independently assayed twice, 52 (16.6%) exhibited a difference in ipTM > 0.1, 19 (6.1%) exhibited a difference in ipTM > 0.2, and 8 (2.5%) exhibited a difference in ipTM > 0.4, despite being identical paired (not pooled) runs.

To test whether the variability we observed (Fig. 2C) is due primarily to false-positive or false-negative errors, we evaluated the AUROC using the mean, maximum, and minimum size-corrected ipTM for the 38,718 protein pairs in multiple pools. Mean size-corrected ipTM was more predictive of real PPIs than either the minimum or the maximum (Table S4), and the same was true for smaller pool sizes (Fig. S7B), suggesting the variability is a combination of false-positive and false-negative errors. Averaging the size-corrected ipTM across multiple replicates modestly improves prediction of known PPI, consistent with what has been observed for earlier versions of AlphaFold (*20*). Together, these data suggest that variability in ipTM scores is intrinsic to AlphaFold3 and includes both false-positive and false-negative interactions.

### Pairwise interactions accurately recapitulate macromolecular complexes

Our strategy for identifying PPIs is inherently pairwise; each protein pair appears in at least one pool. However, *in vivo* protein interactions are rarely that simple: the functional form of many proteins is multimeric (e.g. homodimeric) and many proteins function in the context of macromolecular complexes. This prompts three related questions. First, can protein interactions in large complexes be accurately predicted by considering only pairwise interactions? Second, can structurally homologous but functionally distinct complexes be distinguished? Third, can protein-protein interfaces that are part of a larger complex be correctly predicted? To answer these questions, we considered three well-characterized higher order complexes (the ribosome, ABC transporters, and RNA polymerase) in additional detail. Where applicable, we compared our results to those of a cross-linking mass spectroscopy (XL-MS) study performed with two different cross-linking chemicals (DSS and DSSO, 11.4A and 10.1A crosslinker spacer arm lengths, respectively) in a related organism, *Mycoplasma pneumoniae* (*17*).

The ribosome is a large macromolecular complex (∼2,500 kDa) composed of 2 large rRNA molecules and ∼50 ribosomal proteins. Most ribosomal proteins interact primarily with the rRNA, although some scaffold each other’s interactions. To assess the performance of pooled-AlphaFold3 at predicting these interactions in the absence of the scaffolding 16S and 23S rRNA (which were not included in our predictions), we compared the size-corrected ipTMs of all ribosomal protein interactions with the minimum distances between ribosomal proteins extracted from a recent cryo-EM structure of the *M. pneumoniae* ribosome (*21*). Size-corrected ipTM was largely consistent with the minimum distances extracted from the *M. pneumoniae* ribosome structure (*21*) (Fig. 3A-B). Protein pairs closer than 5 Å exhibited significantly higher ipTMs than those located further apart (Fig. S8; *t*-test *p* = 1.625e-17). Notably, pooled-AlphaFold3 was significantly more accurate at recapitulating protein-protein interactions in the ribosome than XL-MS (*17*) (Fig. 3C, Fig. S8), despite the high expression and stable interactions of these proteins. The ability of pooled-AlphaFold3 prediction to correctly “assemble” the ribosome in the absence of the rRNA scaffold may be due to its efficient use of structural templates (*3*): although the structure (*21*) we used for benchmarking was not part of the default template set used, ribosome structures from other species likely served as accurate templates for ribosomal protein structures and interactions, allowing accurate PPI prediction in the absence of biological context.

**Fig. 3.**
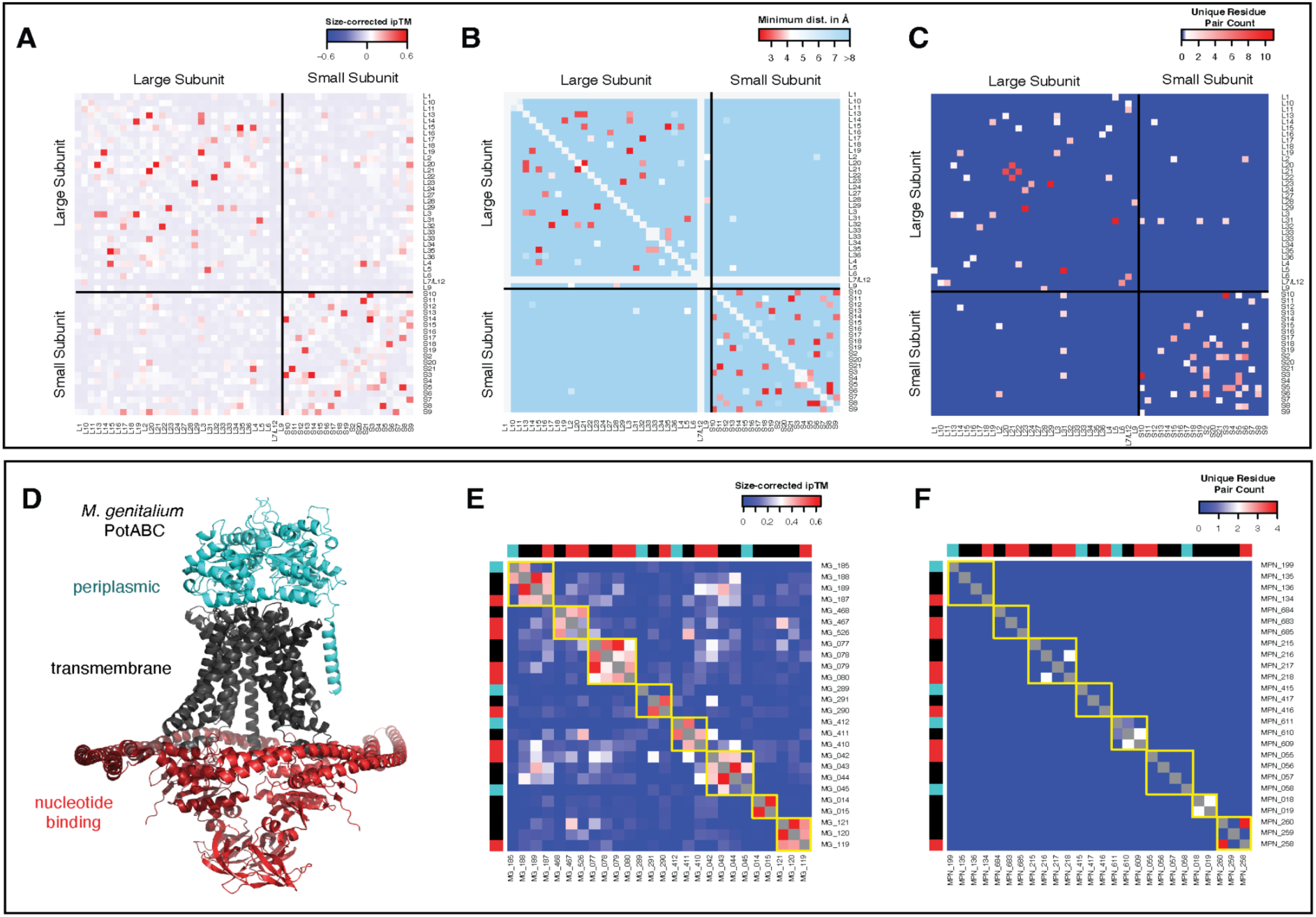
**A.** Size-corrected ipTM values between all ribosomal proteins in *M. genitalium* from the pooled-AlphaFold3 screen. Red denotes high size-corrected ipTM values. Note that almost all high size-corrected ipTM interactions occur between protein pairs in the same ribosomal subunit. **B.** Minimum distance between ribosomal proteins in the published cryo-EM structure of the *M. pneumoniae* ribosome (*21*). Red denotes protein pairs close together. **C.** Unique residue pairs between ribosomal proteins in published *M. pneumoniae* XL-MS data. Red denotes protein pairs with identified cross-links, indicating proximity. For **A**, **B**, and **C** colors are such that nearby protein pairs should be in red, leading to the similar pattern of red dots in all three panels. **D.** Predicted AlphaFold3 complex of the *M. genitalium* ABC transporter PotABC with domains labeled and color-coded. **E.** Size-corrected ipTMs between the eight ABC-type nutrient transporters. **F.** Unique residue pairs between ABC-type transporters in *M. pneumoniae* XL-MS data (*17*). For **E** and **F**, ABC transporter domains (periplasmic, transmembrane, nucleotide-binding) are color coded in the row and column sides consistent with **D**, and interactions within individual ABC transporters are outlined in yellow.

*M. genitalium* encodes eight ABC-type nutrient transporters. ABC transporters are structurally similar (Fig. 3D), consisting of 2 transmembrane (TM) subunits, 2 nucleotide-binding domains (NBD), and (optionally) a periplasmic binding protein (PBP) (*22*), and are frequently encoded in operons. We took advantage of this organization to assess the performance of pooled-AlphaFold3 in correctly assembling a large number of distinct but structurally homologous complexes. For each of the eight ABC transporters, we compared the size-corrected ipTM of biologically relevant interactions (PBP-TM, TM-TM, and TM-NBD) both within a single transporter and between different transporters. All three types of interactions exhibited significantly higher size-corrected ipTMs within a single transporter than between transporters (Fig. 3E), and individual transporters were clearly discernible in an unclustered heatmap of pairwise size-corrected ipTMs, with very little crosstalk (Fig. 3E). Pooled-AlphaFold3 discerned ABC transporters better than XL-MS (Fig. 3F) highlighting the independence of our *in silico* predictions from protein abundance. Our analysis of ABC transporters strongly supports the idea that pooled-AlphaFold3 can sensitively and specifically segregate and assemble structurally homologous complexes.

RNA polymerase is a multisubunit enzyme composed of ααβ’βδσ^A^ and functions as a key regulatory node. We found strong predicted interactions between the 5 subunits of RNA polymerase (RpoABCDE), GreA (a small protein that interacts with the secondary channel to rescue stalled RNA polymerases (*23–25*)), MG_354, a DUF1951 protein previously shown to interact with RNA polymerase (*17*), and other RNA polymerase interacting proteins such as NusG (*26*), UvrD (*27*), and Spx (*28*) (Fig. 4, Table S4). Since many of these interactions have been structurally characterized, we asked whether our largely pairwise interactions accurately recapitulated the structure of interactions in and with this complex. We compared each pairwise structure of RNAP subunits (extracted from pools of other proteins) to a predicted structure of the complete *M. genitalium* holoenzyme (ααβ’βδσ^A^, generated using AlphaFold3; Fig. 4) or experimental structures of relevant complexes (see Table S6).

**Fig. 4.**
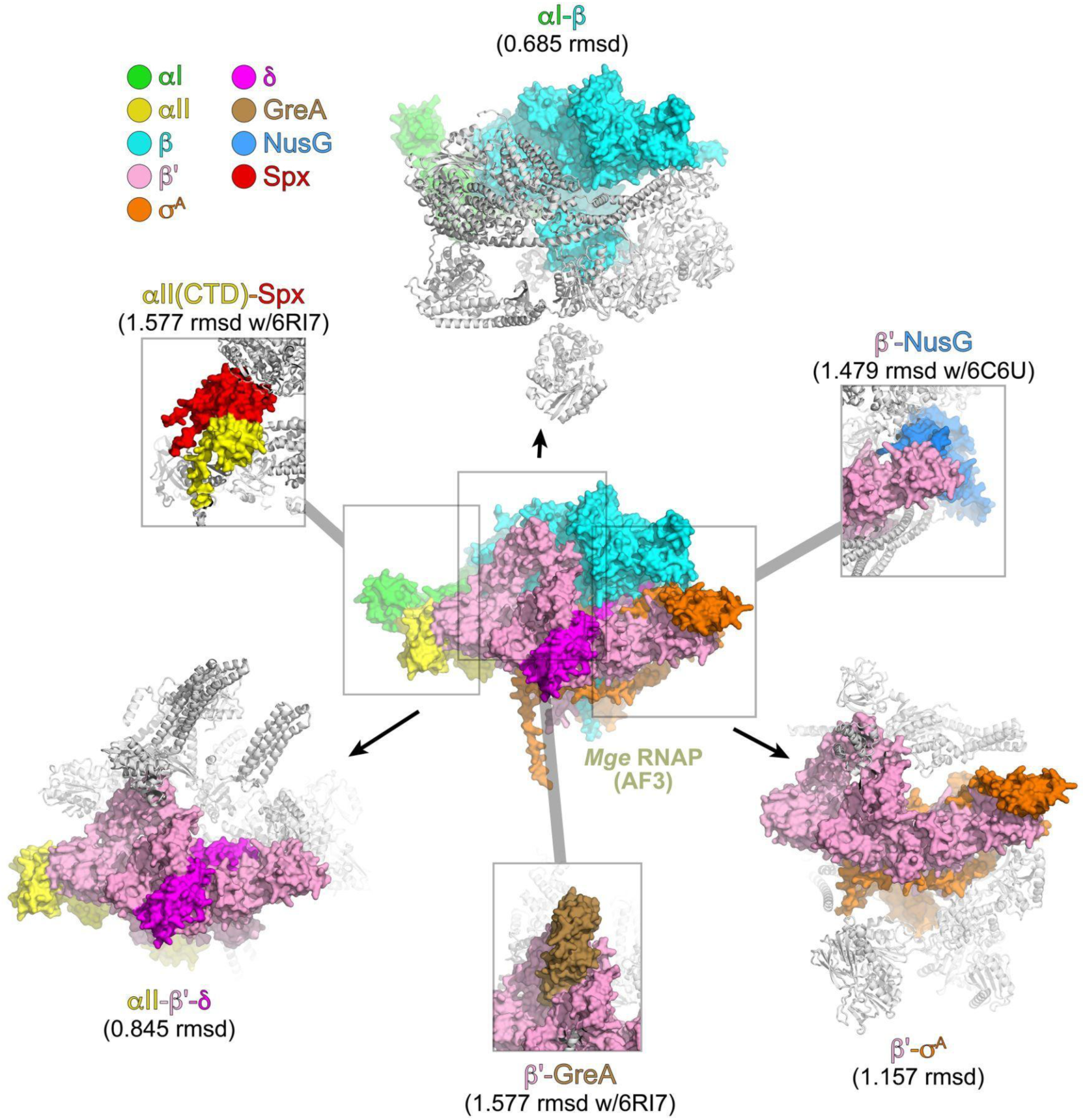
Pooled-AlphaFold3 accurately predicts known interactions with RNAP subunits. Shown in the center is the AF3-predicted structure of *M. genitalium* (Mge) RNAP (αI-αII-β-β’-δ-σ^A^); the proteins are shown as molecular surfaces and color-coded according to the legend (upper left). Shown on the top, lower right, and lower left are selected examples of pooled-AlphaFold3 predictions that included interacting RNAP subunits: (*top*) allbyall1130 included αI (green molecular surface) and β (cyan molecular surface) in a milieu of eight other proteins (shown as gray ribbons). These proteins from the pool superimposed with the equivalent proteins from Mge RNAP with an RMSD of 0.685 Å (Table S6). (*lower right*) allbyall2016 included β’ (pink) and σ^A^ (orange) in a milieu of seven other proteins (gray ribbons). These proteins from the pool superimposed with the equivalent proteins from Mge RNAP with an RMSD of 1.157 Å (Table S6). (*lower left*) allbyall5 included αI (yellow), β’ (pink), and δ (magenta) in a milieu of 14 other proteins (gray ribbons). These proteins from the pool superimposed with the equivalent proteins from Mge RNAP with an RMSD of 0.845 Å (Table S6). Boxed are the PPIs for predictions of other RNAP interacting proteins with RNAP subunits for which experimental structures are available: (*top right*) allbyall1290 included β’ (pink) and NusG (blue) in a milieu of 11 other proteins (gray ribbons). (*bottom*) allbyall1607 included β’ (pink) and GreA (brown) in a milieu of nine other proteins (gray ribbons). (*top left*) allbyall256 included (yellow) and Spx (red) in a milieu of 11 other proteins (gray ribbons).

Pairwise predicted interfaces closely matched the same interfaces in the holoenzyme. For instance, the job allbyall5 co-folded 17 *M. genitalium* proteins, including three RNAP subunits αII, β’, and δ (Table S6), which were correctly predicted to interact (Table S2). The αII-β’-δ complex predicted from allbyall5 superimposed on the same subunits of the *M. genitalium* RNAP with an RMSD of 0.845 Å over 1,091 a-carbons (using the PyMOL align command to exclude flexible loops that do not align well). Predicted interactions between RNAP subunits and other RNAP interacting proteins GreA, NusG, and Spx were compared with available experimental structures (Table S6). Overall,16 jobs included at least one RNAP subunit and another interacting protein (Table S6) and the predicted interactions superimposed with the corresponding proteins in the *M. genitalium* RNAP holoenzyme or with the experimental structures with a mean RMSD of 0.899 ± 0.312 Å (median number of a-carbons included in alignments was 1,109). As with the ribosome, the efficient use of structural templates likely played an important role in the accuracy of these predictions. These data suggest that our inherently pairwise pooled-AlphaFold3 predictions accurately predict not only the interacting partners but also the interaction interfaces (at least for RNA polymerase and other structurally characterized complexes), allowing the use of this method for structural analyses, such as the identification of competitive (or synergistic) protein binding interfaces.

### Pooled-AlphaFold3 suggests novel interactions for M. genitalium proteins

Given that our *M. genitalium* predicted PPIs were highly predictive of known biological interactions and complexes, we visualized the combined ipTM and correlation data (all combined scores > 0.2) as a network (Fig. 5A). Genes of similar function frequently clustered together: transporters, ribosomal proteins, RNA polymerase subunits, and DNA replication/repair proteins each formed dense interconnected networks (Fig. 5A). We used this network to manually assess previously unknown interactions to determine if they could inform new hypotheses about *Mycoplasma* protein functions. Below, we (briefly) highlight three such stories present in our dataset.

**Fig. 5.**
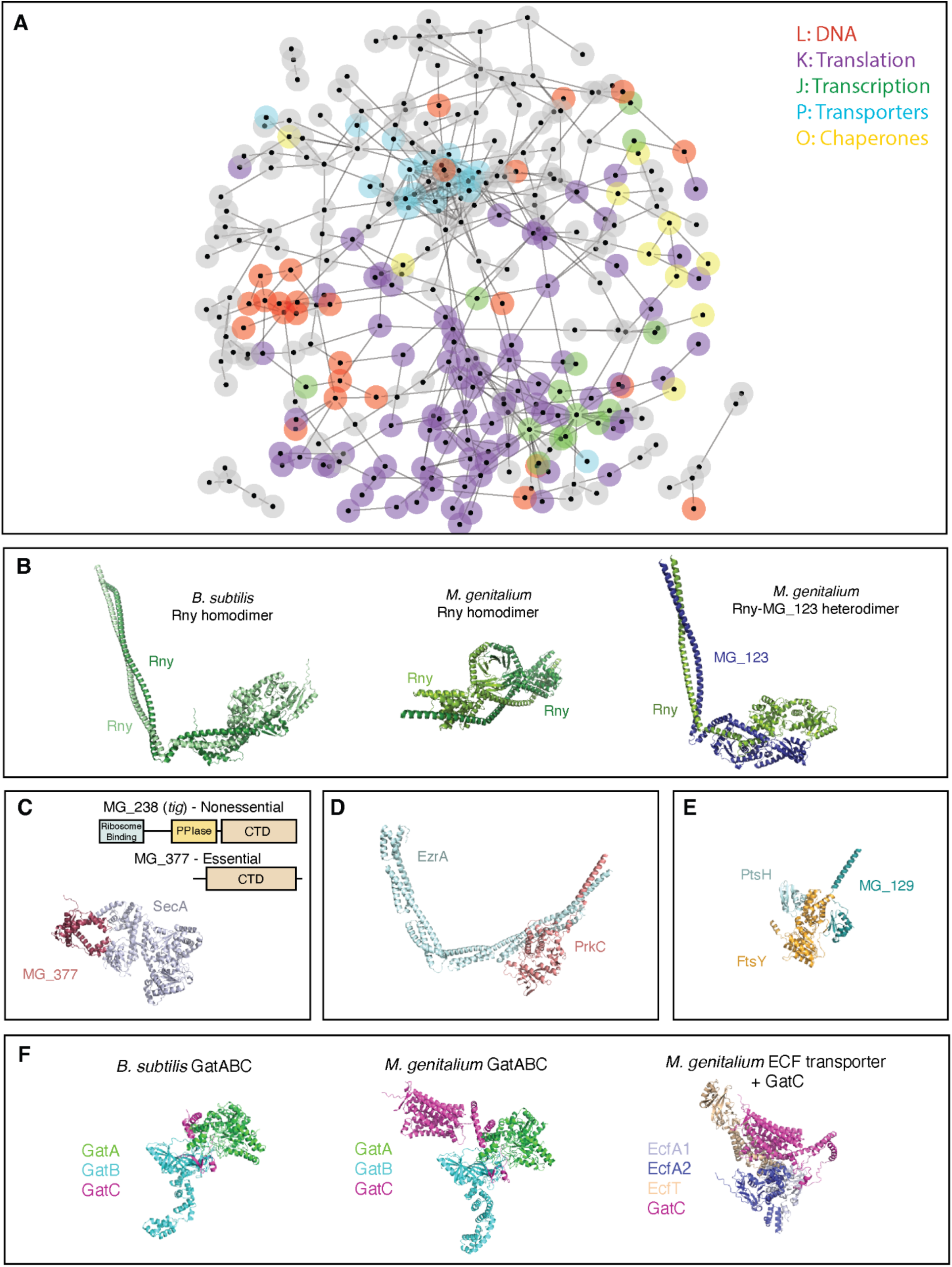
Novel *M. genitalium* predicted PPIs. **A.** Network of genome-wide PPIs in *M. genitalium*. Genes are represented by nodes, and edges are set as (size-corrected ipTM + 0.2 * correlation(ipTM)). Layout is edge-weighted spring-embedded. All edges with weight > 0.2 are shown. Genes in specific COG categories are highlighted by node borders: Translation (J) = purple, Transcription (K) = green, DNA (L) = red, Ion Transport (P) = cyan, Chaperones (O) = yellow. All other COGs, including Unknown (S), are shown in grey; clusters of these proteins may represent spurious interactions. Transcription (RNA polymerase) genes cluster within translation due to the presence of predicted interactions related to transcription-translation coupling (e.g. NusA, NusG, RpsB, RpsE). **B**. Predicted structure of *B. subtilis* Rny homodimer, *M. genitalium* Rny homodimer exhibiting incorrect structure, and *M. genitalium* Rny/MG_123 heterodimer exhibiting correct fold. **C.** Schematic of the domain organization of the two Tig homologs in *M. genitalium* and predicted interaction between SecA and the essential *tig* homolog MG_377. The interacting residues of MG_377 are located near the center of its CTD-domain. **D.** Predicted interaction between EzrA and the serine/threonine kinase PrkC. **E.** Predicted structure of FtsY-PtsH-MG_129 complex, showing independent interfaces of two PTS components with FtsY. **F.** Predicted complexes of *B. subtilis* GatABC, *M. genitalium* GatABC, and *M. genitalium* EcfA_1_A_2_T-GatC, showing the S-protein domain of GatC in *M. genitalium* and its predicted interaction with the ECF transporter.

### A heterodimeric RNase Y in M. genitalium and M. pneumoniae

In Firmicutes, RNase Y is a quasi-essential RNase responsible for mRNA turnover (*29*). Like its functional homolog in gram-negative bacteria (RNase E), RNase Y is membrane tethered and membrane attachment is important for its function (*30*). RNase Y typically forms a homo-dimer, with a coiled-coil domain between the N-terminal transmembrane helix and the catalytic C-terminal forming the principal dimerization interface (Fig. 5B) (*31*). Strikingly, although AlphaFold3 correctly folded the *B. subtilis* Rny homodimer, the *M. genitalium* Rny (MG_130) homodimer folded incorrectly with a low ipTM (Fig. 5B, Table S7). However, these bacteria possess a distant homolog of RNase Y (MG_123/MPN_262, ∼5% identity to RNase Y, member of COG1418). Our pooled-AlphaFold3 data suggest that this protein dimerizes with RNase Y (Fig. 5B), forming a heterodimer instead, and the predicted structure of this complex is consistent with the XL-MS data (7/9 unique crosslinks between Rny and MPN_262 within 30 Å, Table S8). Supporting the idea that RNase Y in these species is a heterodimer, MG_123/MPN_262 is essential, as is RNase Y (*18*). The functional implications of this heterodimeric RNase Y remain to be explored, but may be related to the unusual post-transcriptional regulatory mechanisms postulated for these genome-reduced bacteria (*32*).

### A potential role in protein secretion for an essential truncated trigger factor protein

Trigger factor (Tig) is a ribosomal and cytoplasmic chaperone with an additional role in post-translational protein secretion (*33–35*). Tig consists of three domains: an N-terminal ribosome binding domain, a central FKBP-type peptidyl-prolyl cis-trans isomerase chaperone domain, and a C-terminal domain involved in chaperone functions (Fig. 5C). Trigger factor is ubiquitous in bacterial genomes, non-essential, and almost never duplicated due to a severe dosage constraint caused by the ribosome-binding N-terminal domain (*36*). In the few genomes with multiple trigger factor-like genes, only one usually contains the N-terminal domain (*36*). *M. genitalium* and the related *M. pneumoniae* each encode a full-length non-essential Tig and an essential Tig homolog containing only the C-terminal chaperone domain (MG_377/MPN555, Fig. 5C) (*17*, *18*). Our data suggest a role for MG_377/MPN555 in protein secretion: both *M. genitalium* MG_377 and *M. pneumoniae* MPN555, but not their canonical Tigs, have strong predicted interactions with SecA, the membrane subunit of the bacterial Sec apparatus (Table S4, Table S7, Fig. 5C). This predicted interaction may be essential for SecA function due to the lack of other secretory chaperones such as SecB or CsaA (*37*) in *Mycoplasma*.

### A potential role for serine/threonine kinase in Mycoplasma cell division

Tenericutes including *Mycoplasma* lack a peptidoglycan cell wall, making their cell division process fundamentally different from that of other bacteria, where an FtsZ ring organizes septal peptidoglycan synthesis (*38*). Thus, relatively little is known about cell division in *Mycoplasma.* Our dataset predicts an interaction between the cell division-associated protein EzrA/MG_397 and the eukaryotic-like serine/threonine kinase PrkC/MG_109 (Fig. 5D). AlphaFold3 predicts an interaction between these proteins in both the closely related *M. pneumoniae* and in the more distantly related Tenericute *Spiroplasma citrii* (Table S7), but not between EzrA and any of the three serine/threonine kinases in the distantly related Firmicute *B. subtilis* (Table S7). Deletion of PrkC in *M. pneumoniae* results in a significant decrease in EzrA protein concentration (*39*), a pattern shared with the cytoskeletal HMW1-5 proteins, which require phosphorylation by PrkC for stability in *M. pneumoniae* (*40*). Intriguingly, the spectrin-like domains of EzrA exhibit a coiled-coil motif, as do large portions of the HMW1-5 proteins (*41*) and *B. subtilis* GpsB (a target of PrkC in that species (*42*)), raising the possibility that PrkC preferentially recognizes and phosphorylates coiled-coils to regulate cell division.

### Moonlighting enzymes in essential processes

*M. genitalium* and *M. pneumoniae* are the products of extreme genome reduction, which has been associated with neofunctionalization of proteins leading to moonlighting enzymes (*43*). Below, we discuss two novel examples of moonlighting revealed in our data.

FtsY is involved in SRP-mediated co-translational protein targeting to the membrane (*44*). We were surprised to find that our pooled-AlphaFold3 screen identified two proteins involved in the phosphoenolpyruvate-dependent sugar phosphotransferase system (PTS) as interactors of FtsY: MG_129, an orphan PTS EIIB protein (size corrected ipTM = 0.496), and MG_041, the phosphocarrier protein PtsH (size corrected ipTM = 0.317). A single crosslink between FtsY and MPN_268/MG_129 was identified in the XL-MS study (*17*), which connected residues 33.4 Å apart in the predicted complex (Fig. 5E, Table S8). No single EIIC or EIIA components are encoded in the *M. genitalium* genome, supporting the idea that MG_129 moonlights in a different process, such as co-translational protein secretion.

ECF-transporters consist of EcfA1, EcfA2, and EcfT components which combine with a specificity protein (S) to allow the import of various micronutrients (*45*). Strikingly, both GatC and GatB, components of the essential Glu-tRNA^Gln^ amidotransferase (GatABC (*46*)), were predicted to interact with ECF transporter components. The heterotrimeric enzyme GatABC transamidates mis-acylated Glu-tRNA^Gln^, thereby functionally replacing glutaminyl-tRNA synthetase (which is not found in gram-positive bacteria such as *M. genitalium*). Analysis of the predicted structure of *M. genitalium* GatC revealed an additional ∼300aa N-terminal domain with structural homology to S-proteins which is not present in GatC proteins outside of Mycoplasma and Ureaplasma and is predicted to bind to the same spot on the ECF transporter as an S-protein (Fig. 5F). The interaction between GatC and the ECF transporter is supported by the *M. pneumoniae* XL-MS data (*17*), which identifies cross-links between GatC and several components of the ECF transporter consistent with our predicted structure (3/3 crosslinks between GatC and ECF transporter components within 30 Å; Table S8). The functional implications of this interaction remain to be explored: the S-protein domain of GatC may moonlight in the import of a substrate for its transamidase function, may import a completely unrelated metabolite, or may affect GatABC function via membrane anchoring.

## Discussion

Genome-scale PPI prediction is a powerful strategy for revealing new biology, but its broad adoption has been hampered by its high computational demands and a significant false positive rate. Pooled-PPI prediction increases the accuracy of genome-scale approaches while decreasing the overall runtime as well as the number of discrete jobs of such screens and can even allow some genome-scale queries to be run on free web-based resources, democratizing access to this powerful tool. We demonstrate the utility of this strategy by predicting all pairwise PPIs (113,050) in the *M. genitalium* genome using only 2,027 jobs, which required just 68 person-days using our personal AlphaFold3 accounts. Our results reveal a wealth of technical insights into AlphaFold3 as well as biological insights into *M. genitalium* RNA metabolism, protein secretion, and cell division that will drive future experimental studies.

Our study assessed all possible pairwise interactions in an organism (rather than carefully curated subsets of known negative and positive interactions) and included replicates for ∼1/3 of them (by default), allowing us to uncover several important features of AlphaFold3. First, our study revealed that pooled-AlphaFold3 significantly outperforms paired AlphaFold3 for identifying known PPIs, by largely eliminating false-positive interactions through a “decoy” mechanism. Second, the preponderance of non-interacting protein pairs in our data enabled us to visualize and correct for the size dependence of ipTM scores. Although several shortcomings of ipTM scores have been previously described (*47–49*), the size dependence of non-interacting ipTM scores has not been explicitly noted. Correcting for this bias greatly increased the performance of ipTM in predicting known interactions and is likely to be applicable to other studies as well. Third, our dataset revealed the variability of AlphaFold3 ipTM scores in both pooled and paired settings. This variability has likely remained unappreciated both because the size bias of ipTM scores obfuscates discrepancies and because jobs are seldom replicated due to the high computational costs. Finally, our in-depth analysis of predicted PPIs between ribosomal proteins, RNA-polymerase subunits, and ABC-transporters revealed that AlphaFold3 does not generally require fully-functional biological complexes to accurately predict interacting proteins and their interfaces, or to distinguish structurally homologous (but functionally distinct) complexes.

Pooled-PPI prediction usefully identifies known and biologically relevant PPIs, and can (in limited cases) enable the use of web-interfaces to perform genome-scale PPI prediction. It can be used to query all pairwise interactions between a set of proteins (all-by-all), or (less efficiently) for few-by-all screens. Facile access to highly specific genome-scale PPI screens has major implications. First, combining genome-wide PPI information with high-dimensional phenotypic screens, such as chemical genomics, can uncover mechanistic insights into functional protein-protein interactions, greatly accelerating our understanding of biological processes by highlighting novel connections that may not be credible in the absence of orthogonal data. Second, future pooled-PPI prediction screens could also incorporate RNAs and other molecules to assay their interacting partners: such studies could uncover novel tRNA-, rRNA- and/or ncRNA-interacting proteins. Third, genome-scale PPI screens can be used to probe the biology of uncultured organisms, such as the Asgard archaea, which contain the relatives of the last eukaryotic common ancestor. Finally, PPI screens can be used to study intractable or fleeting interactions between organisms, such as between hosts and pathogens, members of ecological communities, or communities and hosts (e.g. the microbiome), potentially revealing novel protein determinants and drug targets. Pooled-PPI prediction democratizes the ability to make genome-scale PPI predictions (whether all-by-all or selected subsets), allowing more labs to apply this tool to more problems and ultimately furthering our understanding of biology.

## Acknowledgments

We thank Google DeepMind and Isomorphic Labs for providing free online access to AlphaFold3 to us and to the greater research community. We thank T. Kortemme, T. Goddard, D. Todor, J. Rock, D. Booth, A. Johnson, M. Garber, S. Silas, A. Typas, and members of the Gross and Beltrao Labs for extensive helpful discussions.

## Author contributions

Conceptualization: HT, LMK, CAG, PB Methodology: HT, LMK, JJ, SAD, PB, CAG Investigation: HT, LMK, JJ, HNB, SAD Visualization: HT, LMK, JJ, SAD

Funding acquisition: CAG, PB, SAD Project administration: HT, CAG Supervision: HT, PB, CAG

Writing – original draft: HT, LMK, JJ, HNB, SAD, PB, CAG Writing – review & editing: HT, LMK, JJ, HNB, SAD, PB, CAG

## Data and materials availability

All data derived from alphafoldserver.com was deposited under the original AlphaFold3 Server Output License in zenodo: https://doi.org/10.5281/zenodo.15499631

All locally generated AlphaFold3 data was deposited under the AlphaFold3 license in zenodo: https://zenodo.org/records/16920556

## Supplementary Materials

Materials and Methods

Supplementary Text

Figs. S1 to S8

Tables S1 to S7

## METHODS

### Running AlphaFold3

All 2,027 comprehensive pools as well as some paired jobs were run using default settings on the AlphaFold3 server (https://alphafoldserver.com/). Additional paired and pooled jobs were run locally using AlphaFold 3.0.1 and default settings.

### Analysis of AlphaFold3

All AlphaFold3 runs (pooled or pairs) were downloaded. Raw ipTM scores were extracted from the “summary_confidences” files for all 5 models and averaged. The ipTM of protein pairs that appeared in multiple pools was averaged across all pools unless otherwise noted.

### Preparing pools

The genome of *M. genitalium* was downloaded from NCBI (GenBank: L43967.2, Table S1). Proteins were pooled to minimize the number of pools required to co-fold all protein pairs at least once while keeping all jobs <5,000 tokens. For the 2k, 3k, 4k, and 5k pool samples, the genome was divided into 5 approximately equal sections and proteins from each section were randomly pooled to cover all possible pairs.

### Size correction of ipTM scores

To determine the effect of summed protein size on ipTM, we performed a robust linear regression using the robustbase::lmrob function, which computes a MM-type regression estimator (a robust and efficient estimator with a ∼50% breakdown point and 95% efficiency). A robust estimator was used to mitigate the influence of true-positive interactions on the regression. The linear regression was performed on the square-root of the summed size of the two proteins (in amino acids).

### STRING data

STRING data related to taxid: 243273 was downloaded on 3/18/2025 from https://string-db.org/. Only the experimental channel was used for benchmarking PPIs.

### Structural analysis

For all of the structural analyses, we focused on the top scoring model. The analysis of RNAP is described in Table S6. For the ribosome, mmCIF files were loaded into R and manipulated using the Bio3D package (*50*). Distances and RMSD were computed using the functions in this package. The structure of the *M. pneumoniae* ribosome (*21*) (7P6Z) downloaded from PDB was used.

### Crosslinking analysis

XL-MS data (*17*) was mapped onto AlphaFold3 models of relevant *M. pneumoniae* protein complexes using PyXlinkViewer (*51*). Satisfaction threshold was set to 30 Å. Both “internal” and “between” crosslinks were mapped for all proteins within the complex.

## SUPPLEMENTARY FIGURES

**Figure S1.**
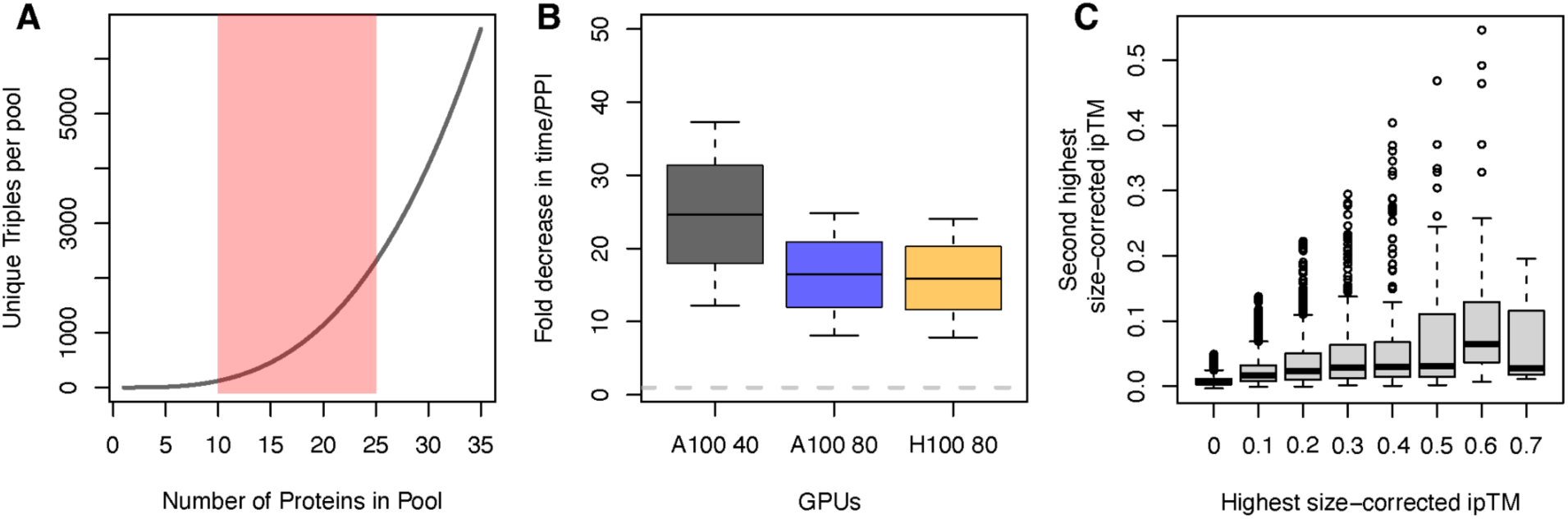
Pooled-PPI prediction drastically reduces the number of required jobs and increases inference throughput for identifying tripartite interactions. **A.** The number of proteins in the pool determines the number of unique tripartite interactions screened in a single job. The red shaded area encompasses most jobs. **B.** Pooled-PPI-prediction decreases inference time per unique tripartite PPI even considering the increased runtime of larger jobs. All timing estimates (1024-5120 tokens) and pool sizes of 10-25 proteins are considered and reflected in the error bars. **C.** Tripartite interactions are rare in our data set – most proteins in most jobs have 0 or 1 strong interactions. Even in cases where a protein strongly interacts with 2 partners, identifying a true tripartite interaction would require having additional pools with each of the two partners individually and most large complexes are accurately modeled in a pairwise manner (described later in the manuscript).

**Fig. S2.**
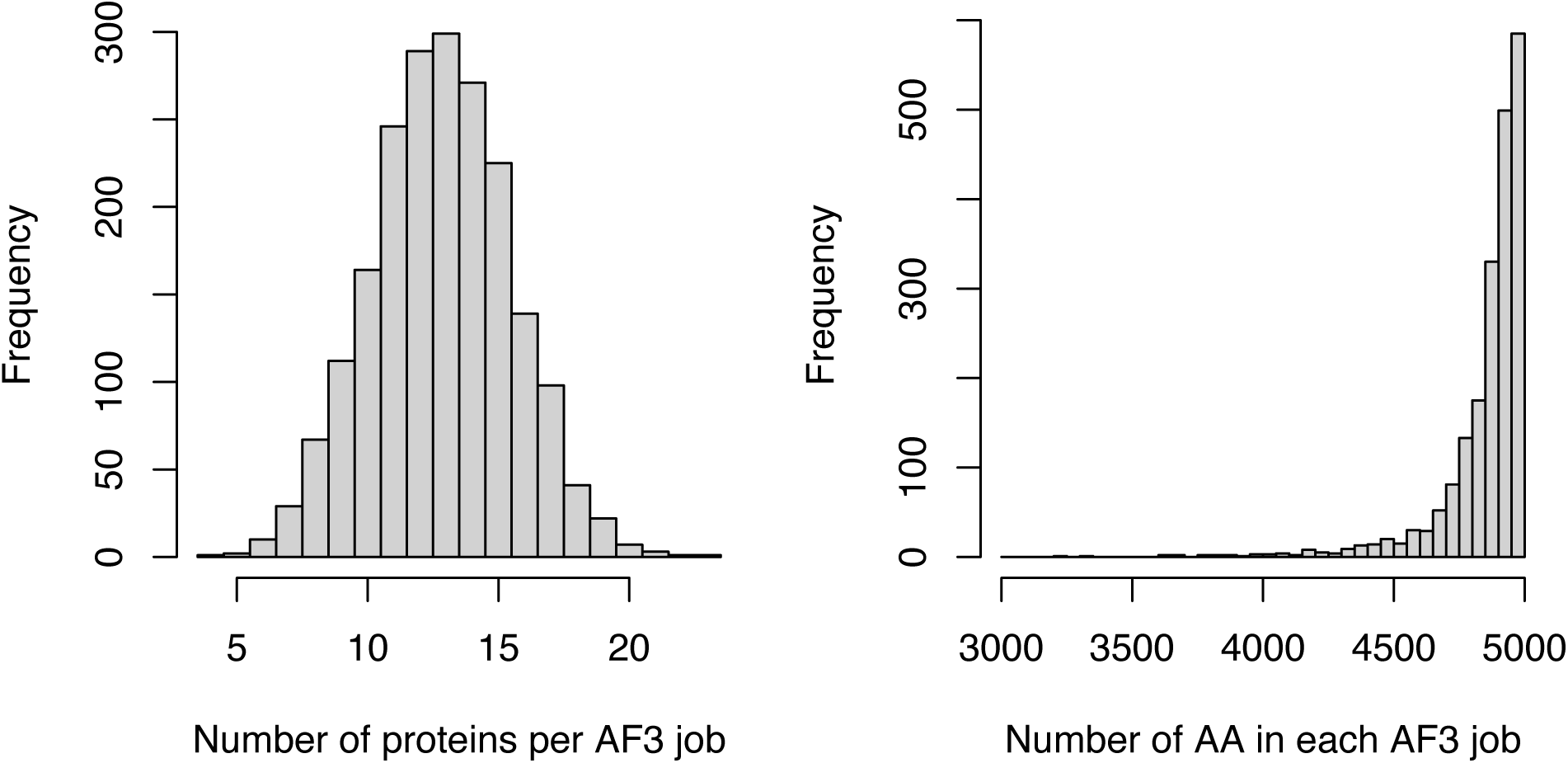
Characteristics of the *M. genitalium* pools. **A.** Histogram of pool sizes (Table S1 and S2). **B.** Histogram of total job size (in amino acids, Table S1 and S2).

**Fig. S3.**
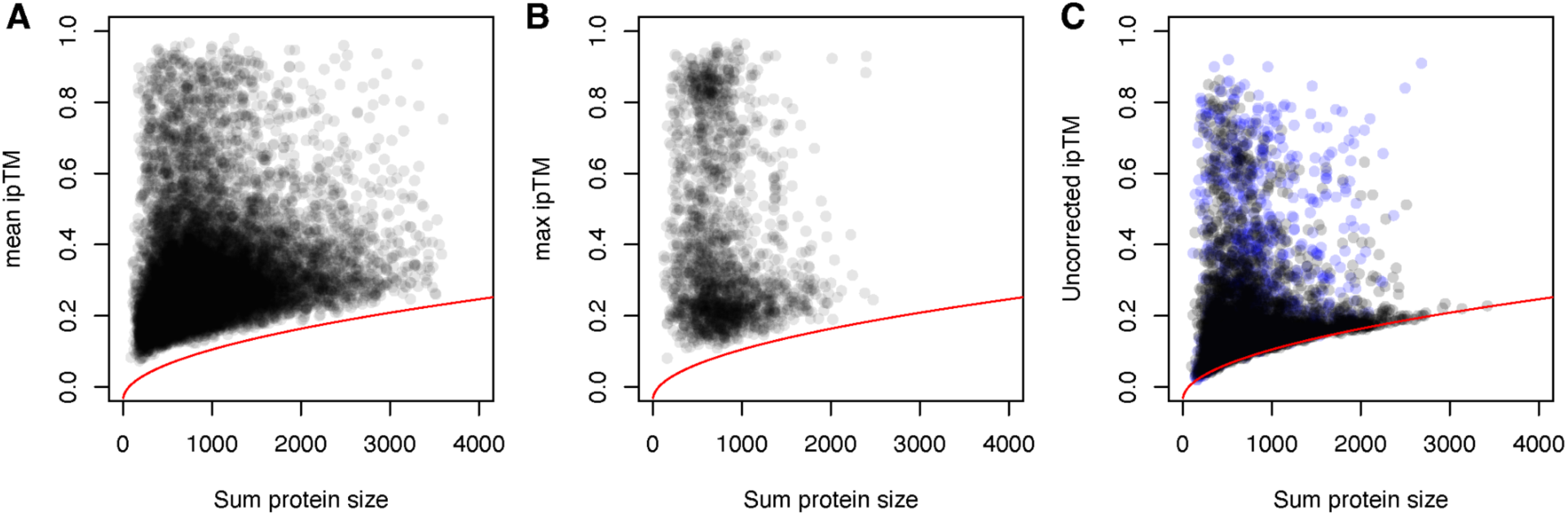
Size-bias of ipTM scores is not limited to pooled approaches or to AlphaFold3. Red line in all panels is the fit line from AlphaFold3 pools: sqrt(sum_protein_size)*0.0044 - 0.036. **A.** Summed protein size plotted by ipTM for AlphaFold-Multimer data from ref (*4*) exhibits a similar pattern to our data, albeit a lower correlation (robust R^2^ = 0.162, n = 13,274), likely due to a lower proportion of non-interacting protein pairs. **B.** Summed protein size plotted by ipTM for AlphaFold-Multimer data from ref (*52*) exhibits a similar pattern to our data, albeit an even lower correlation (robust R^2^ = 0.021, n = 1,977), likely due to a very low proportion of non-interacting protein pairs. **C.** Summed protein size plotted by ipTM for individual pairs folded by AlphaFold3 exhibits a similar pattern albeit a lower correlation. Two sets of data are shown. (1) Protein pairs selected primarily from high-scoring interactions in pools and run on alphafoldserver.com (robust R^2^ = 0.170, n = 942, blue). (2) Random sample of 4,560 protein pairs run using a local implementation of AlphaFold3 (robust R^2^ = 0.236, n = 4,560,black).

**Figure S4.**
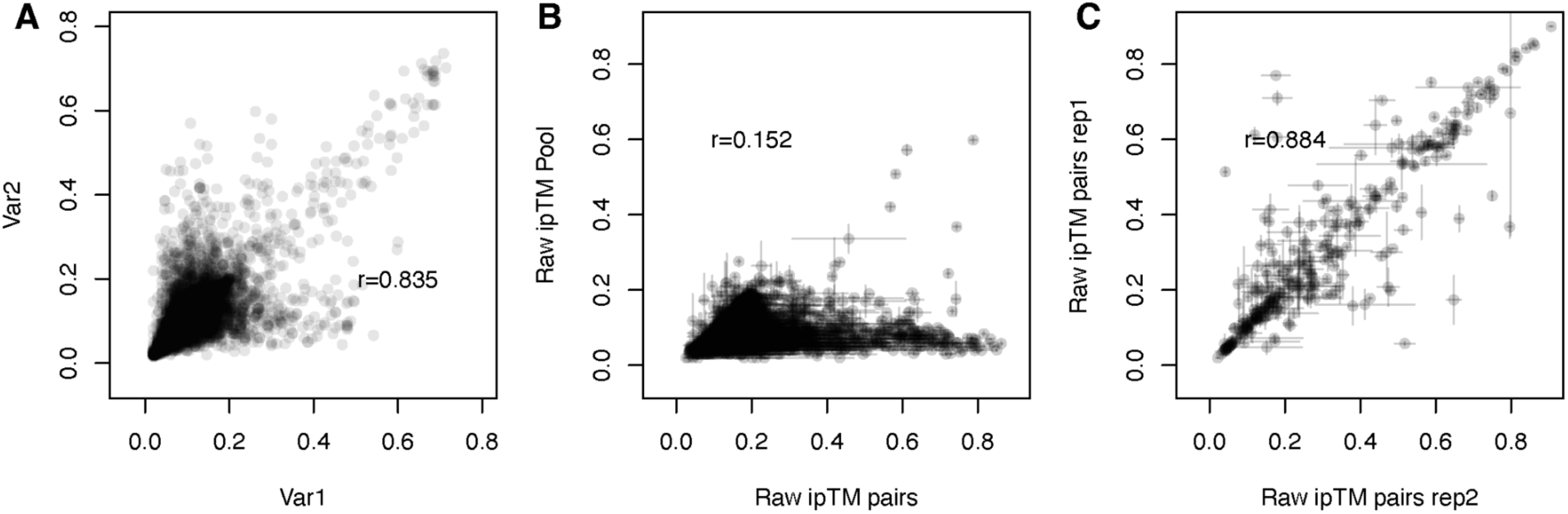
Uncorrected ipTM values make replicates look better than they are. **A.** Raw ipTM scores for the 38,718 protein pairs that appear in multiple pools are similar (Pearson’s *r* = 0.835, 38,718 protein pairs). **B.** Raw ipTM is similar in paired and pooled AlphaFold3 jobs (Pearson’s *r* = 0.152, 4,560 pairs). **C.** Raw ipTM scores across replicated paired runs exhibit surprising variability (Pearson’s *r* = 0.884, 314 pairs).

**Fig. S5.**
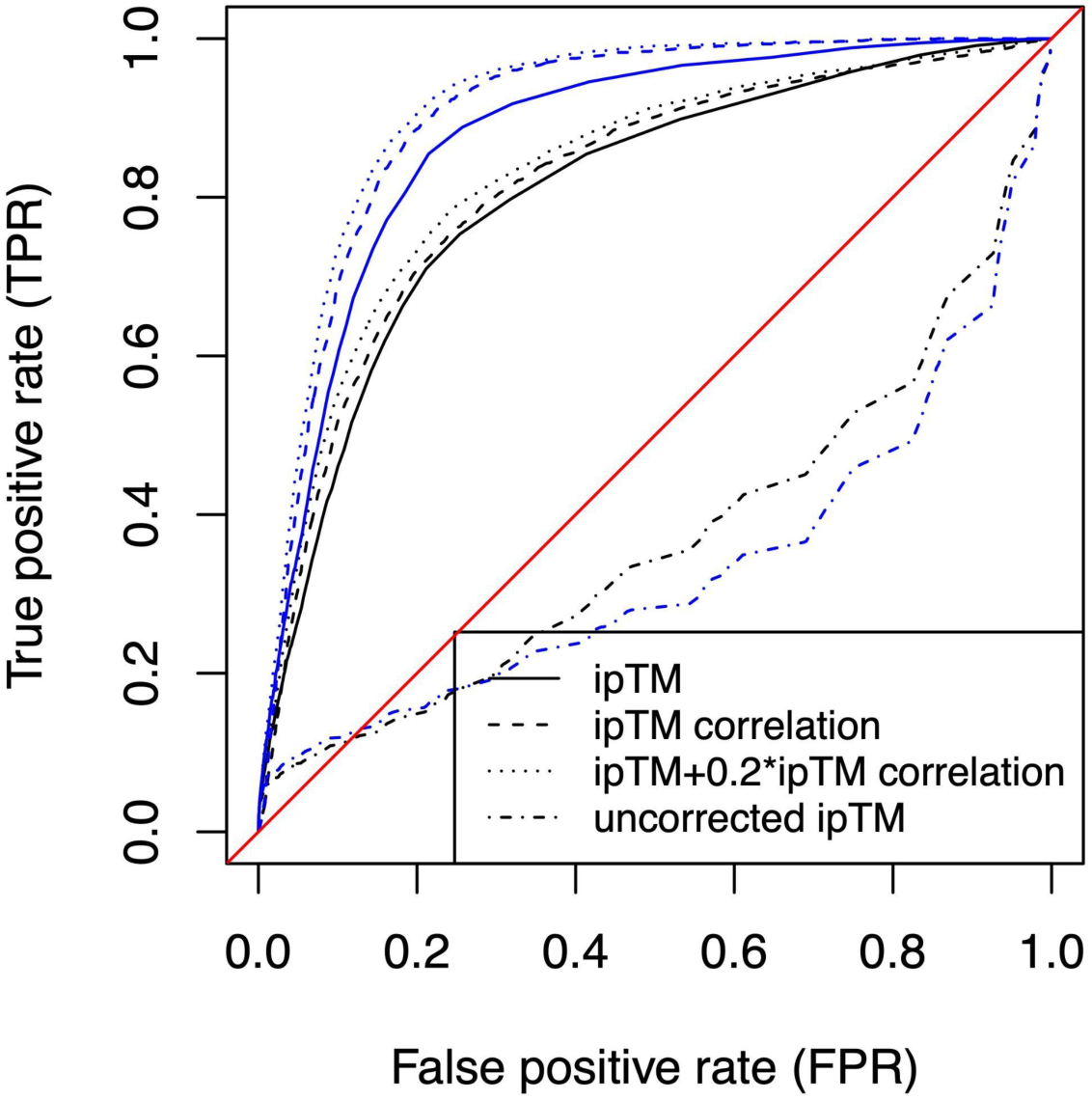
Pooled-AlphaFold3 accurately predicts known interactions in the STRING database. AUROC curve for size-corrected ipTM scores (solid line), size-corrected ipTM correlations (large dashed line), size-corrected ipTM + 0.2 size-corrected ipTM correlation (small dashed line), and uncorrected ipTM (alternating small and large dashed line). Black lines consider interactions with STRING experimental scores > 800 (strong interactions) as the true positive set. Blue lines consider interactions with STRING experimental scores = 999 (strongest interactions) as the true positive set.

**Figure S6.**
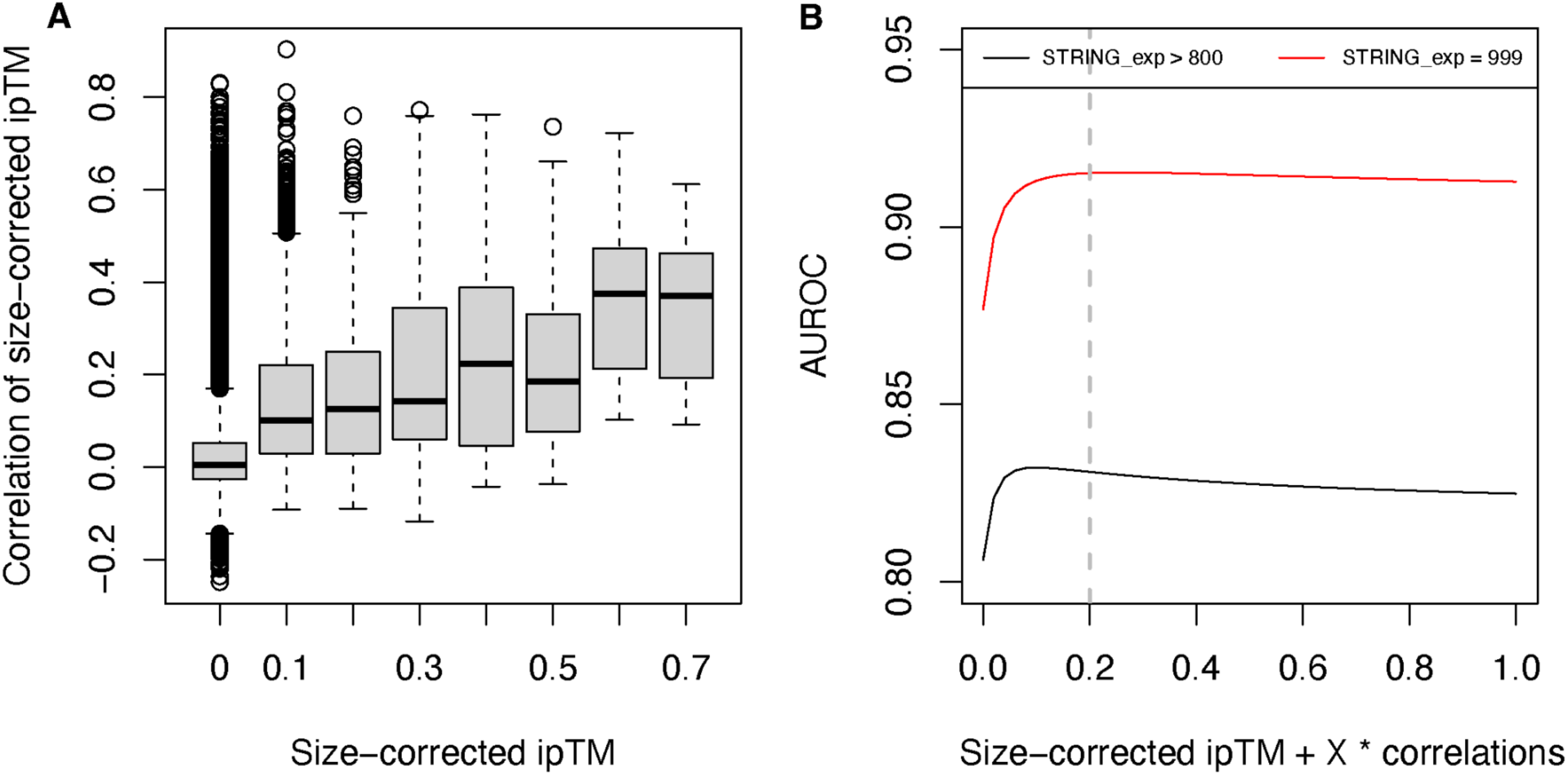
Size-corrected ipTM and the correlation between the size-corrected ipTM of two proteins are partially orthogonal, and combining the two increases predictive performance. **A.** Boxplot showing partial correlation between the size-corrected ipTM of two proteins and the correlation between their size-corrected ipTM profiles. **B.** AUROC benchmarking the STRING experimental dataset with different combinations of size-corrected ipTM and its correlation. For STRING > 800 (black line), the maximum AUROC is 0.832 and is achieved at (size-corrected ipTM + 0.1 correlation). For STRING = 999 (red line), the maximum AUROC is 0.915 and is achieved at (size-corrected ipTM + 0.26 correlation). We use a combined score of (size-corrected ipTM + 0.2 correlation) for the remainder of the manuscript (dashed vertical line).

**Fig. S7.**
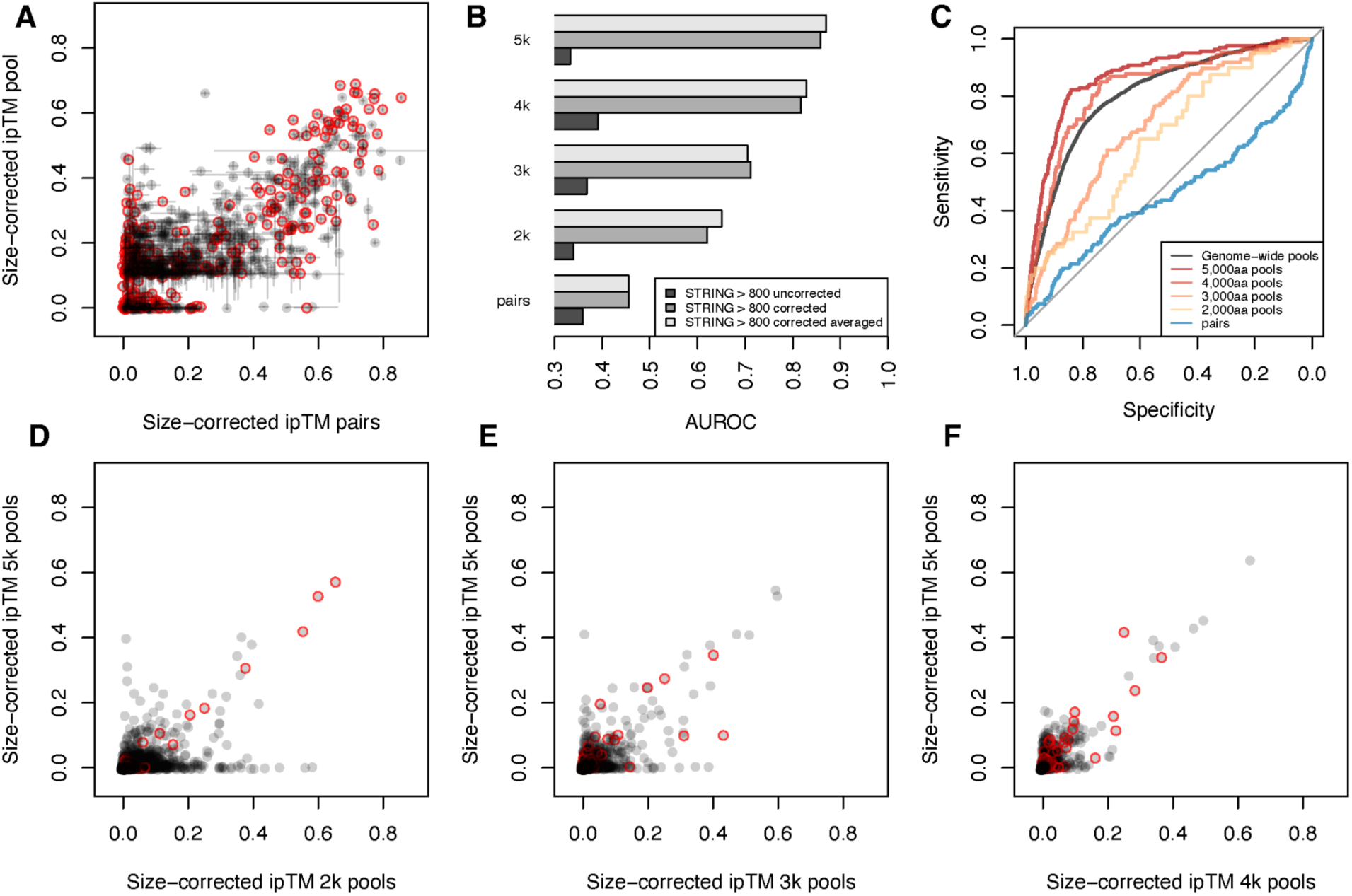
Larger pool sizes are more predictive of known PPIs and exhibit fewer false positive hits. **A.** Size-corrected ipTMs of protein pairs with high scores in the pools are well correlated. (Pearson’s r = 0.725, 942 pairs), though ∼0 to 0.2 lower in pools. **B.** AUROC for ∼4,500 protein pairs assayed using randomly generated pools of different sizes. Raw, corrected, and corrected averaged data is shown. **C.** ROC curves for the different pool sizes. **D-F.** Comprehensive pooled ipTMs compared to ipTMs from 2,000aa (**D**), 3,000aa (**E**), and 4,000aa (**F**) pools. Red circles in **A** and **D-F** represent protein pairs with STRING experimental scores > 800.

**Fig. S8.**
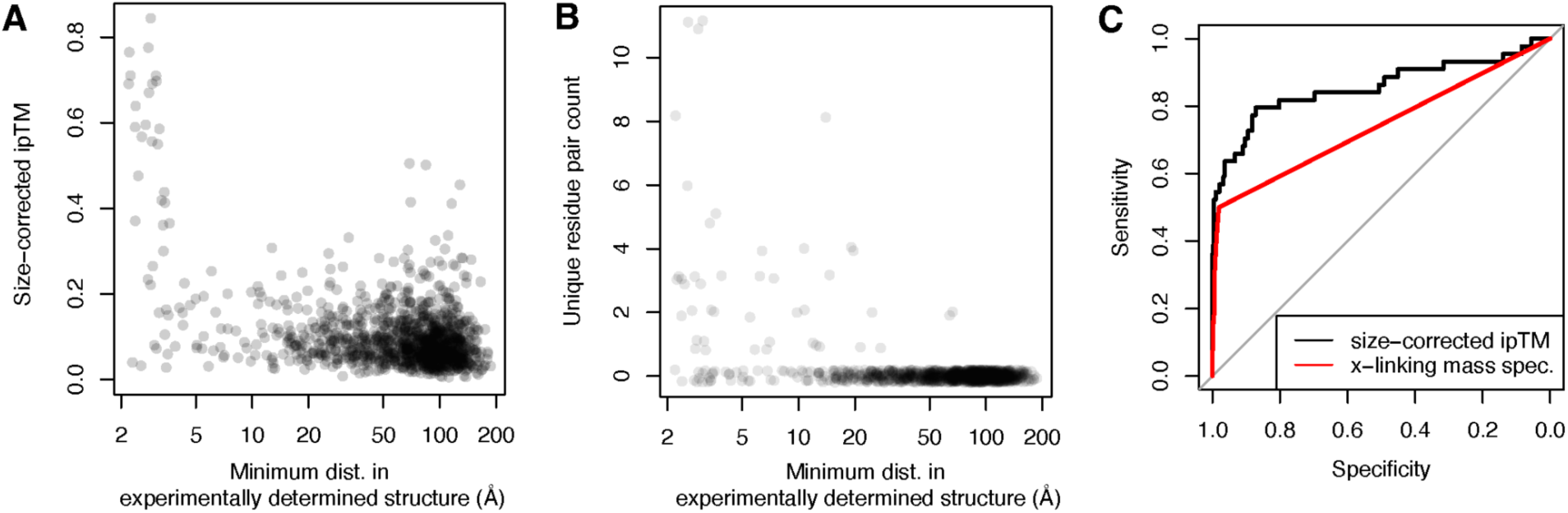
Additional information about the ribosome structure prediction. **A.** Plot of size-corrected ipTM versus minimum distance for all ribosomal protein pairs. **B.** Plot of number of unique crosslinks versus minimum distance for all ribosomal protein pairs. Some of the additional crosslinks in the XL-MS data may be due to the linker length of the cross-linking reagents. **C.** AUROC curve showing the performance of XL-MS (red) and size-corrected ipTM for identifying ribosomal protein pairs within 5 Å.

## SUPPLEMENTARY TEXT 1

Raw ipTM scores demonstrated a strong and significant correlation with the square root of the summed size of interacting proteins. This relationship allows us to calculate a “predicted-ipTM” for each protein pair. There are two straightforward ways to correct ipTM scores:

(1) (actual_ipTM - predicted_ipTM)
(2) (actual_ipTM / predicted_ipTM)

In the manuscript, we use the subtraction-based approach “size-corrected ipTM” (1), because its similarity to raw ipTM makes existing intuitions broadly applicable. However, the division- based approach “ipTM-ratio” (2) actually performs better than size-corrected ipTM in predicting known interactions (Fig. N1, Table S4).

**Fig. N1.**
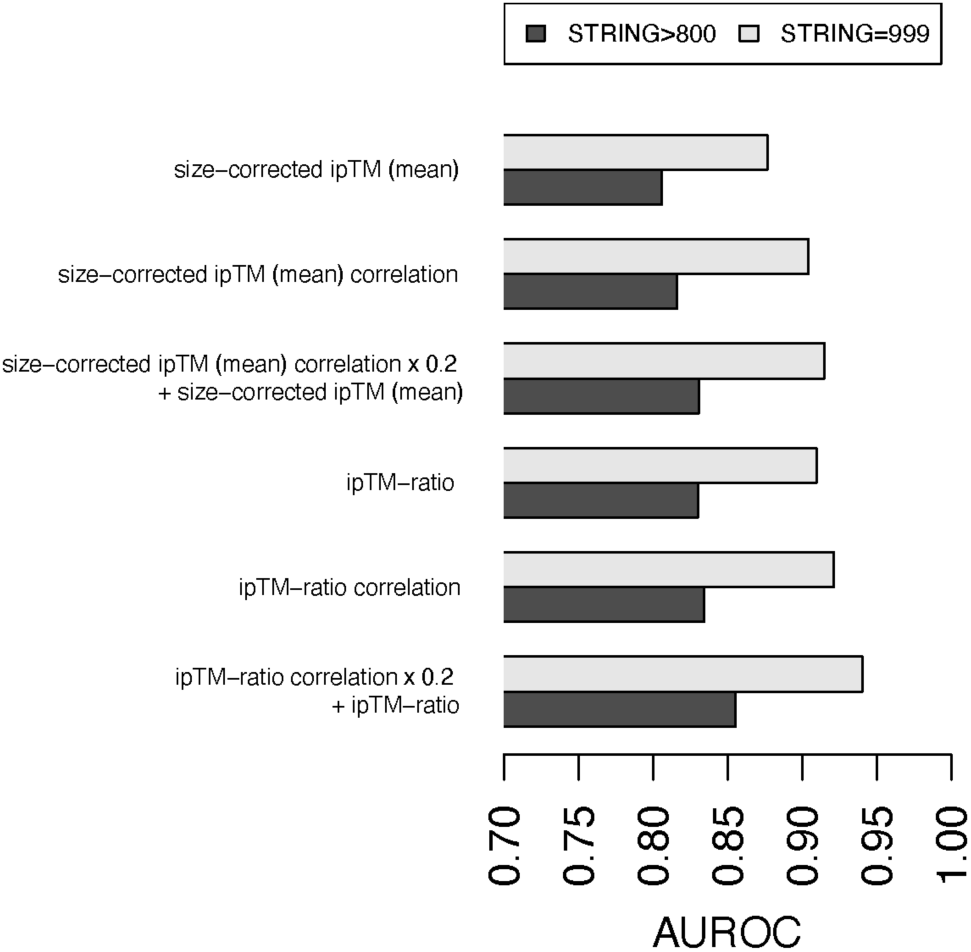
Barplot showing AUROC values for different methods of correcting size bias. Data is in Table S4.

The success of a division-based size correction in predicting known interactions suggests that interactions involving small proteins (<200aa) are significantly undervalued by both raw ipTM and the subtraction-based size-corrected ipTM used in this manuscript. For these proteins, ipTMs within the “borderline” range of 0.05-0.2 using the subtraction-based correction may represent strong interactions, warranting close attention. ipTM-ratio values are included in Table S4.

## References

1. J. Jumper, R. Evans, A. Pritzel, T. Green, M. Figurnov, O. Ronneberger, K. Tunyasuvunakool, R. Bates, A. Žídek, A. Potapenko, A. Bridgland, C. Meyer, S. A. A. Kohl, A. J. Ballard, A. Cowie, B. Romera-Paredes, S. Nikolov, R. Jain, J. Adler, T. Back, S. Petersen, D. Reiman, E. Clancy, M. Zielinski, M. Steinegger, M. Pacholska, T. Berghammer, S. Bodenstein, D. Silver, O. Vinyals, A. W. Senior, K. Kavukcuoglu, P. Kohli, D. Hassabis, Highly accurate protein structure prediction with AlphaFold. Nature 596, 583– 589 (2021).

2. R. Evans, M. O’Neill, A. Pritzel, N. Antropova, A. Senior, T. Green, A. Žídek, R. Bates, S. Blackwell, J. Yim, O. Ronneberger, S. Bodenstein, M. Zielinski, A. Bridgland, A. Potapenko, A. Cowie, K. Tunyasuvunakool, R. Jain, E. Clancy, P. Kohli, J. Jumper, D. Hassabis, Protein complex prediction with AlphaFold-Multimer, Bioinformatics (2021). https://www.biorxiv.org/content/10.1101/2021.10.04.463034v2.

3. J. Abramson, J. Adler, J. Dunger, R. Evans, T. Green, A. Pritzel, O. Ronneberger, L. Willmore, A. J. Ballard, J. Bambrick, S. W. Bodenstein, D. A. Evans, C.-C. Hung, M. O’Neill, D. Reiman, K. Tunyasuvunakool, Z. Wu, A. Žemgulytė, E. Arvaniti, C. Beattie, O. Bertolli, A. Bridgland, A. Cherepanov, M. Congreve, A. I. Cowen-Rivers, A. Cowie, M. Figurnov, F. B. Fuchs, H. Gladman, R. Jain, Y. A. Khan, C. M. R. Low, K. Perlin, A. Potapenko, P. Savy, S. Singh, A. Stecula, A. Thillaisundaram, C. Tong, S. Yakneen, E. D. Zhong, M. Zielinski, A. Žídek, V. Bapst, P. Kohli, M. Jaderberg, D. Hassabis, J. M. Jumper, Accurate structure prediction of biomolecular interactions with AlphaFold 3. Nature 630, 493–500 (2024).

4. E. W. Schmid, J. C. Walter, Predictomes, a classifier-curated database of AlphaFold-modeled protein-protein interactions. Mol. Cell 85, 1216–1232.e5 (2025).

5. D. Yu, G. Chojnowski, M. Rosenthal, J. Kosinski, AlphaPulldown-a python package for protein-protein interaction screens using AlphaFold-Multimer. Bioinformatics 39 (2023).

6. E. Yirmiya, S. J. Hobbs, A. Leavitt, I. Osterman, C. Avraham, D. Hochhauser, B. Madhala, M. Skovorodka, J. M. J. Tan, H. C. Toyoda, I. Chebotar, M. Itkin, S. Malitsky, G. Amitai, P. J. Kranzusch, R. Sorek, Structure-guided discovery of viral proteins that inhibit host immunity. Cell 188, 1681–1692.e17 (2025).

7. D. F. Burke, P. Bryant, I. Barrio-Hernandez, D. Memon, G. Pozzati, A. Shenoy, W. Zhu, A. S. Dunham, P. Albanese, A. Keller, R. A. Scheltema, J. E. Bruce, A. Leitner, P. Kundrotas, P. Beltrao, A. Elofsson, Towards a structurally resolved human protein interaction network. Nat. Struct. Mol. Biol. 30, 216–225 (2023).

8. H. Schweke, M. Pacesa, T. Levin, C. A. Goverde, P. Kumar, Y. Duhoo, L. J. Dornfeld, B. Dubreuil, S. Georgeon, S. Ovchinnikov, D. N. Woolfson, B. E. Correia, S. Dey, E. D. Levy, An atlas of protein homo-oligomerization across domains of life. Cell 187, 999–1010.e15 (2024).

9. C. Elfmann, J. Stülke, Cutting-edge tools for structural biology: bringing AlphaFold to the people. Trends Microbiol. 0 (2025).

10. K. Venkatesan, J.-F. Rual, A. Vazquez, U. Stelzl, I. Lemmens, T. Hirozane-Kishikawa, T. Hao, M. Zenkner, X. Xin, K.-I. Goh, M. A. Yildirim, N. Simonis, K. Heinzmann, F. Gebreab, J. M. Sahalie, S. Cevik, C. Simon, A.-S. de Smet, E. Dann, A. Smolyar, A. Vinayagam, H. Yu, D. Szeto, H. Borick, A. Dricot, N. Klitgord, R. R. Murray, C. Lin, M. Lalowski, J. Timm, K. Rau, C. Boone, P. Braun, M. E. Cusick, F. P. Roth, D. E. Hill, J. Tavernier, E. E. Wanker, A.-L. Barabási, M. Vidal, An empirical framework for binary interactome mapping. Nat. Methods 6, 83–90 (2009).

11. L. Chang, A. Perez, Ranking peptide binders by affinity with AlphaFold. Angew. Chem. Int. Ed Engl. 62, e202213362 (2023).

12. P. Vosbein, P. Paredes Vergara, D. T. Huang, A. R. Thomson, AlphaFold ensemble competition screens enable peptide binder design with single-residue sensitivity. ACS Chem. Biol. 19, 2198–2205 (2024).

13. A. Mondal, B. Singh, R. H. Felkner, A. De Falco, G. Swapna, G. T. Montelione, M. J. Roth, A. Perez, A computational pipeline for accurate prioritization of protein-protein binding candidates in high-throughput protein libraries. Angew. Chem. Int. Ed Engl. 63, e202405767 (2024).

14. 14. Docs/performance.md at Main · Google-deepmind/alphafold3 (Github; https://github.com/google-deepmind/alphafold3/blob/main/docs/performance.md).

15. S. Viz-Lasheras, A. Gómez-Carballa, X. Bello, I. Rivero-Calle, A. I. Dacosta, M. Kaforou, D. Habgood-Coote, A. J. Cunnington, M. Emonts, J. A. Herberg, V. J. Wright, E. D. Carrol, S. C. Paulus, W. Zenz, D. S. Kohlfürst, N. Schweintzger, M. Van der Flier, R. de Groot, L. J. Schlapbach, P. Agyeman, A. J. Pollard, C. Fink, T. T. Kuijpers, S. Anderson, U. Von Both, M. Pokorn, D. Zavadska, M. Tsolia, H. A. Moll, C. Vermont, M. Levin, F. Martinón-Torres, A. Salas, EUCLIDS, PERFORM, and DIAMONDS consortia, A diagnostic host-specific transcriptome response for Mycoplasma pneumoniae pneumonia to guide pediatric patient treatment. Nat. Commun. 16, 673 (2025).

16. D. G. Gibson, G. A. Benders, C. Andrews-Pfannkoch, E. A. Denisova, H. Baden-Tillson, J. Zaveri, T. B. Stockwell, A. Brownley, D. W. Thomas, M. A. Algire, C. Merryman, L. Young, V. N. Noskov, J. I. Glass, J. C. Venter, C. A. Hutchison 3rd, H. O. Smith, Complete chemical synthesis, assembly, and cloning of a Mycoplasma genitalium genome. Science 319, 1215–1220 (2008).

17. F. J. O’Reilly, L. Xue, A. Graziadei, L. Sinn, S. Lenz, D. Tegunov, C. Blötz, N. Singh, W. J. H. Hagen, P. Cramer, J. Stülke, J. Mahamid, J. Rappsilber, In-cell architecture of an actively transcribing-translating expressome. Science 369, 554–557 (2020).

18. J. I. Glass, N. Assad-Garcia, N. Alperovich, S. Yooseph, M. R. Lewis, M. Maruf, C. A. Hutchison 3rd, H. O. Smith, J. C. Venter, Essential genes of a minimal bacterium. Proc. Natl. Acad. Sci. U. S. A. 103, 425–430 (2006).

19. D. Szklarczyk, R. Kirsch, M. Koutrouli, K. Nastou, F. Mehryary, R. Hachilif, A. L. Gable, T. Fang, N. T. Doncheva, S. Pyysalo, P. Bork, L. J. Jensen, C. von Mering, The STRING database in 2023: protein-protein association networks and functional enrichment analyses for any sequenced genome of interest. Nucleic Acids Res. 51, D638–D646 (2023).

20. B. Wallner, AFsample: improving multimer prediction with AlphaFold using massive sampling. Bioinformatics 39 (2023).

21. L. Xue, S. Lenz, M. Zimmermann-Kogadeeva, D. Tegunov, P. Cramer, P. Bork, J. Rappsilber, J. Mahamid, Visualizing translation dynamics at atomic detail inside a bacterial cell. Nature 610, 205–211 (2022).

22. A. L. Davidson, J. Chen, ATP-binding cassette transporters in bacteria. Annu. Rev. Biochem. 73, 241–268 (2004).

23. S. Borukhov, A. Polyakov, V. Nikiforov, A. Goldfarb, GreA protein: a transcription elongation factor from Escherichia coli. Proc. Natl. Acad. Sci. U. S. A. 89, 8899–8902 (1992).

24. N. Opalka, M. Chlenov, P. Chacon, W. J. Rice, W. Wriggers, S. A. Darst, Structure and function of the transcription elongation factor GreB bound to bacterial RNA polymerase. Cell 114, 335–345 (2003).

25. M. ’men Abdelkareem, C. Saint-André, M. Takacs, G. Papai, C. Crucifix, X. Guo, J. Ortiz, A. Weixlbaumer, Structural basis of transcription: RNA polymerase backtracking and its reactivation. Mol. Cell 75, 298–309.e4 (2019).

26. J. Y. Kang, R. A. Mooney, Y. Nedialkov, J. Saba, T. V. Mishanina, I. Artsimovitch, R. Landick, S. A. Darst, Structural basis for transcript elongation control by NusG family universal regulators. Cell 173, 1650–1662.e14 (2018).

27. A. A. Kawale, B. M. Burmann, UvrD helicase-RNA polymerase interactions are governed by UvrD’s carboxy-terminal Tudor domain. *Commun*. Biol. 3, 607 (2020).

28. J. Shi, F. Li, A. Wen, L. Yu, L. Wang, F. Wang, Y. Jin, S. Jin, Y. Feng, W. Lin, Structural basis of transcription activation by the global regulator Spx. Nucleic Acids Res. 49, 10756– 10769 (2021).

29. M. Benda, S. Woelfel, P. Faßhauer, K. Gunka, S. Klumpp, A. Poehlein, D. Kálalová, H. Šanderová, R. Daniel, L. Krásný, J. Stülke, Quasi-essentiality of RNase Y in Bacillus subtilis is caused by its critical role in the control of mRNA homeostasis. Nucleic Acids Res. 49, 7088–7102 (2021).

30. S. Laalami, M. Cavaiuolo, J. Oberto, H. Putzer, Membrane localization of RNase Y is important for global gene expression in Bacillus subtilis. Int. J. Mol. Sci. 25, 8537 (2024).

31. N. Morellet, P. Hardouin, N. Assrir, C. van Heijenoort, B. Golinelli-Pimpaneau, Structural insights into the dimeric form of Bacillus subtilis RNase Y using NMR and AlphaFold. Biomolecules 12, 1798 (2022).

32. E. Yus, V. Lloréns-Rico, S. Martínez, C. Gallo, H. Eilers, C. Blötz, J. Stülke, M. Lluch-Senar, L. Serrano, Determination of the gene regulatory network of a genome-reduced bacterium highlights alternative regulation independent of transcription factors. Cell Syst. 9, 143–158.e13 (2019).

33. J. De Geyter, A. G. Portaliou, B. Srinivasu, S. Krishnamurthy, A. Economou, S. Karamanou, Trigger factor is a bona fide secretory pathway chaperone that interacts with SecB and the translocase. EMBO Rep. 21, e49054 (2020).

34. E. Martinez-Hackert, W. A. Hendrickson, Promiscuous substrate recognition in folding and assembly activities of the trigger factor chaperone. Cell 138, 923–934 (2009).

35. K. Wu, T. C. Minshull, S. E. Radford, A. N. Calabrese, J. C. A. Bardwell, Trigger factor both holds and folds its client proteins. Nat. Commun. 13, 4126 (2022).

36. T. Wan, L. Zhuo, Z. Pan, R.-Y. Chen, H. Ma, Y. Cao, J. Wang, J.-J. Wang, W.-F. Hu, Y.-J. Lai, M. Hayat, Y.-Z. Li, Dosage constraint of the ribosome-associated molecular chaperone drives the evolution and fates of its duplicates in bacteria. MBio 15, e0199424 (2024).

37. D. Linde, R. Volkmer-Engert, S. Schreiber, J. P. Müller, Interaction of the Bacillus subtilis chaperone CsaA with the secretory protein YvaY. FEMS Microbiol. Lett. 226, 93–100 (2003).

38. M. Lluch-Senar, E. Querol, J. Piñol, Cell division in a minimal bacterium in the absence of ftsZ: ftsZ and cell division in M. genitalium. Mol. Microbiol. 78, 278–289 (2010).

39. V. van Noort, J. Seebacher, S. Bader, S. Mohammed, I. Vonkova, M. J. Betts, S. Kühner, R. Kumar, T. Maier, M. O’Flaherty, V. Rybin, A. Schmeisky, E. Yus, J. Stülke, L. Serrano, R. B. Russell, A. J. R. Heck, P. Bork, A.-C. Gavin, Cross-talk between phosphorylation and lysine acetylation in a genome-reduced bacterium. Mol. Syst. Biol. 8, 571 (2012).

40. S. R. Schmidl, K. Gronau, C. Hames, J. Busse, D. Becher, M. Hecker, J. Stülke, The stability of cytadherence proteins in Mycoplasma pneumoniae requires activity of the protein kinase PrkC. Infect. Immun. 78, 184–192 (2010).

41. M. F. Balish, Mycoplasma pneumoniae, an underutilized model for bacterial cell biology. J. Bacteriol. 196, 3675–3682 (2014).

42. F. Pompeo, E. Foulquier, B. Serrano, C. Grangeasse, A. Galinier, Phosphorylation of the cell division protein GpsB regulates PrkC kinase activity through a negative feedback loop in Bacillus subtilis: Regulation of PrkC activity by GpsB. Mol. Microbiol. 97, 139–150 (2015).

43. Y. D. Kelkar, H. Ochman, Genome reduction promotes increase in protein functional complexity in bacteria. Genetics 193, 303–307 (2013).

44. J. D. Miller, H. D. Bernstein, P. Walter, Interaction of E. coli Ffh/4.5S ribonucleoprotein and FtsY mimics that of mammalian signal recognition particle and its receptor. Nature 367, 657–659 (1994).

45. S. Rempel, W. K. Stanek, D. J. Slotboom, ECF-type ATP-binding cassette transporters. Annu. Rev. Biochem. 88, 551–576 (2019).

46. K. Sheppard, D. Söll, On the evolution of the tRNA-dependent amidotransferases, GatCAB and GatDE. J. Mol. Biol. 377, 831–844 (2008).

47. A.-R. Kim, Y. Hu, A. Comjean, J. Rodiger, S. E. Mohr, N. Perrimon, Enhanced Protein-Protein Interaction Discovery via AlphaFold-Multimer, Bioinformatics (2024). https://www.biorxiv.org/content/10.1101/2024.02.19.580970v1.

48. R. L. Dunbrack Jr, Rēs ipSAE loquunt : What’s wrong with AlphaFold’s ipTM score and how to fix it, Bioinformatics (2025). https://www.biorxiv.org/content/10.1101/2025.02.10.637595v1.

49. J. K. Varga, S. Ovchinnikov, O. Schueler-Furman, actifpTM: a refined confidence metric of AlphaFold2 predictions involving flexible regions. Bioinformatics 41, btaf107 (2025).

50. B. J. Grant, L. Skjaerven, X.-Q. Yao, The Bio3D packages for structural bioinformatics. Protein Sci. 30, 20–30 (2021).

51. B. Schiffrin, S. E. Radford, D. J. Brockwell, A. N. Calabrese, PyXlinkViewer: A flexible tool for visualization of protein chemical crosslinking data within the PyMOL molecular graphics system. Protein Sci. 29, 1851–1857 (2020).

52. F. J. O’Reilly, A. Graziadei, C. Forbrig, R. Bremenkamp, K. Charles, S. Lenz, C. Elfmann, L. Fischer, J. Stülke, J. Rappsilber, Protein complexes in cells by AI-assisted structural proteomics. Mol. Syst. Biol. 19, e11544 (2023).

